# One Health or Three? Transmission modelling of *Klebsiella* isolates reveals ecological barriers to transmission between humans, animals and the environment

**DOI:** 10.1101/2021.08.05.455249

**Authors:** Harry Thorpe, Ross Booton, Teemu Kallonen, Marjorie J. Gibbon, Natacha Couto, Virginie Passet, Juan Sebastian Lopez Fernandez, Carla Rodrigues, Louise Matthews, Sonia Mitchell, Richard Reeve, Sophia David, Cristina Merla, Marta Corbella, Carolina Ferrari, Francesco Comandatore, Piero Marone, Sylvain Brisse, Davide Sassera, Jukka Corander, Edward J. Feil

## Abstract

The *Klebsiella* group is highly diverse both genetically and ecologically, being commonly recovered from humans, livestock, plants, soil, water, and wild animals. Many species are opportunistic pathogens, and can harbour diverse classes of antimicrobial resistance (AMR) genes. *K. pneumoniae* is responsible for a high public-health burden, due in part to the rapid spread of health-care associated clones that are non-susceptible to carbapenems. *Klebsiella* thus represents a highly pertinent taxon for assessing the risk to public health posed by animal and environmental reservoirs. Here we report an analysis of 6548 samples and 3,482 genome sequences representing 15 *Klebsiella* species sampled over a 15-month period from a wide range of clinical, community, animal and environmental settings in and around the city of Pavia, in the northern Italian region of Lombardy. Despite carbapenem-resistant clones circulating at a high frequency in the hospitals, we find no genotypic or phenotypic evidence for non-susceptibility to carbapenems outside of the clinical environment. The non-random distribution of species and strains across sources point to ecological barriers that are likely to limit AMR transmission. Although we find evidence for occasional transmission between settings, hierarchical modelling and intervention analysis suggests that direct transmission from the multiple non-human (animal and environmental) sources included in our sample accounts for less than 1% of hospital disease, with the vast majority of clinical cases originating from other humans.

## Introduction

The *Klebsiella* genus is a member of the *Enterobacteriacae* family, related to other enteric pathogens such as *Salmonella* and *E. coli*. By far the best studied *Klebsiella* species is *K. pneumoniae*, which the WHO has recognised as a critical-priority health-care associated pathogen^1^. Antibiotic resistance has spread rapidly within *K. pneumoniae*, and other members of the genus, at least since the early 1980s^2^, and over the last two decades the emergence and spread of genes encoding carbapenemases, that confer non-susceptibility to carbapenems, is of particular concern^3,4^. These genes are most commonly carried on plasmids, and fall into five major groups: *bla*_OXA-48_, *bla*_KPC_, *bla*_VIM_, *bla*_NDM_ and *bla*_IMP_. Widespread clones of *K. pneumoniae* and other *Klebsiella* species that are associated with these genes are primarily spread through the health-care network^5^. In addition, and in common with other key AMR determinants, genes encoding carbapenemases have been reported in multiple non-clinical settings including livestock and wastewater^6–8^.

Increasing concern that these environmental reservoirs of antibiotic resistance pose direct and indirect risk to public health has led to a widening adoption of the One-Health framework for AMR management^9^. This integrative approach is underpinned by a synthesis of antibiotic stewardship and AMR surveillance within clinical, community, agricultural and environmental settings. However, existing data on the abundance and distribution of AMR strains and genes in the environment does not provide a full picture, and our current understanding of their maintenance and spread within and between ecological settings remains fragmentary^10^. Given the urgent requirement for policy priorities informed by robust risk assessments, this represents a key knowledge gap.

A powerful approach to inferring pathogen transmission dynamics is to use whole genome sequencing (WGS) on focal taxa, combined with phylogenetic and clustering analyses, and other statistical methods. Although WGS has been applied successfully to bacterial pathogens within health-care settings^11–14^, and has played a key role in the management of the SARS-CoV-2 pandemic^15^, capturing transmission pathways within ‘*the environment*’ presents significant challenges, and requires large contemporaneous samples from well-defined geographic regions. Nevertheless, a picture has begun to emerge that the risk of transmission of AMR genes and strains from environmental or agricultural settings into the clinic may be rather low, at least in well-resourced regions; a view that is at odds with the prevailing One-Health *cri de coeur*^*16*^. Whole genome sequencing has been used to argue that transmission of AMR strains and/or genes between humans and agricultural animals is limited in *E. coli*^*17*^, *Enterococcus faecium*^*18*^ and *K. pneumoniae*^*19,20*^. The evidence, however, remains equivocal^21^; a compelling counter-example is the study on colistin resistance dissemination in humans in China, which was shown to be largely driven by aquaculture activities^22^.

The sequencing of large and carefully sampled collections of isolates holds the promise to inform an overarching model describing the rate of transmission between settings^23^, and to simultaneously shed light on the relevant biological and ecological factors underpinning transmission barriers. Whilst commonalities of gene and community profiles between settings point to ample opportunities for mixing, the risks to public health of environmental reservoirs of AMR remain difficult to gauge in the absence of this broad framework ^10,24,25^. Recent advances in bioinformatics tools and analytical approaches provide the means to extract critical added value from genome data, and thus to provide a more nuanced view of how microbes and mobile elements move through complex ecosystems.

Here we report a large-scale One-Health study based on 6,548 samples and WGS data for 3,482 isolates encompassing 15 *Klebsiella* (including *Raoutella* species^26^) approximately half of which are *K. pneumoniae*. These data were generated in order to identify environmental reservoirs that pose the biggest risks to public health by quantifying the frequency of transmission between clinical and non-clinical settings. Samples from multiple clinical, community, veterinary, agricultural and environmental sources were taken within a 15-month period around a single city, Pavia, in northern Italy. This represents an unprecedented contemporaneous sampling and sequencing effort within a restricted geographical area that is a known hotspot for health-care associated multiply resistant *K. pneumoniae*^*27*^. We describe the distribution of species, strains, AMR and virulence genes in different settings, and evidence from an intervention model to quantify the impact of transmission between animal and environmental sources, and health-care settings. In addition, these data shed light on the phylogeny and diversity of the *Klebsiella* genus, including the identification of novel lineages of potential species status, and provide reference data for future investigation of the population structures of individual species.

## Materials and methods

### Sampling

Samples were collected in the city of Pavia (Northern Italy) and the surrounding province between July 2017 and October 2018. Information on the 6548 samples collected are given in Table S1. To summarise, the following types of samples were collected: stool and rectal swabs from hospital inpatients and outpatients (four different hospitals) and from a nursing home; stool from healthy volunteers; stool and rectal swabs from companion animals, farm animals or animals admitted in veterinary clinics (dogs, cats, cattle, pigs, poultry, turtles, rabbits and wild birds); invertebrates; samples of edible and ornamental plants, both wild and purchased from groceries, garden centres and large-scale distribution; soil samples; samples of drinking water (drinking fountains) and surface water (rivers and irrigation ditches); surface swabs from hospital, anthropic surfaces (including ATM keypads, ticket machines, buses, benches, supermarket trolleys) and farm surfaces (including enclosure, buckets, milking machines). *Klebsiella* isolates obtained from the laboratory diagnostic routine from urine, wound swabs, respiratory samples and blood cultures of hospital patients with infections were also processed.

### Sample Processing

Stool and rectal swab samples (Fecal swabs, Copan, Brescia, Italy), both from human and animals, were enriched in Luria Bertani (LB) broth with amoxicillin (10 mg/ml) at 36±1°C for 24 hours. Invertebrates were frozen for at least 24 hours after sampling, and surviving bacteria were recovered from the surface of the animals as well as the gut. For the surface, each insect was washed with sterile water for 2 minutes and an aliquot of the washing was enriched in LB broth with amoxicillin (10 mg/ml). For the gut, the insect’s surface was washed with ethanol 70% for 5 minutes and then air-dried. Small insects were ground with a pestle, while larger ones were dissected with sterile scalpel to separate the gut. The gut was then used to inoculate LB broth with amoxicillin (10 mg/ml), which was left to incubate at 36±1°C for 24 hours.

For plants, each sample was divided into four portions: rhizosphere, rhizoplane, epiphyte, endophyte. The portions corresponding to the rhizosphere, rhizoplane and epiphytes were washed with PBS 1X (pH 7.2). The buffer used for washing was then added to LB broth with amoxicillin (10 mg/ml) at 36±1°C for 24 hours. Endophytes were washed with ethanol 70% for 2 minutes and rinsed with sterile water before being washed with a sodium hypochlorite 2% and Triton X-100 1% solution and incubated for 2 minutes before washing with sterile water. Endophytes were ground in PBS 1X (pH 7.2) with pestle and mortar. An aliquot of 1 ml was enriched in LB broth with amoxicillin (10 mg/ml) at 36±1°C for 24 hours.

Soil samples (5 grams) were washed in PBS 1X (pH 7.2), which was then added to LB broth with amoxicillin (10 mg/ml) at 36±1°C for 24 hours. Water samples (1L for both drinking and river water) were filtered through a sterile filter unit (pore size 0.45µm, 0.2 µm; Thermo Scientific) and the membranes were enriched in LB broth with amoxicillin (10 mg/ml) at 36±1°C for 24 hours. For environmental water (mainly from ditches and ponds) we sampled and filtered at least 50 ml of water (higher volumes when possible) and then proceeded in the same way as the drinking and river waters. Surface swabbing was performed on areas of 10 cm^2^ for each point by using a swab rinse kit (Copan, Brescia, Italy). After the collection, the swab and its medium were enriched in LB broth with amoxicillin (10 mg/ml) at 36±1 °C for 24 hours.

For each of the above samples, one microliter of each enrichment was plated on Simmons Citrate Agar with Inositol (SCAI)^28,29^ medium and the plates were incubated at 36±1°C for 48 hours.

### Species identification and antimicrobial susceptibility testing

Yellow and mucoid colonies on SCAI plates suspected to belong to the *Klebsiella* genus were identified at the species level through MALDI-TOF MS (Microflex LT/SH Bruker Daltonik GmbH, Bremen, Germany) equipped with Bruker biotyper 3.1 software (Microflex LT/SH Bruker Daltonik GmbH). Once confirmed to be members of this genus, the isolates were subcultured on MacConkey agar for antibiotic susceptibility testing and DNA extraction. Antibiotic susceptibility was tested for all the isolates using the BD Phoenix 100 automated system (Boston, Dickinson and Company, Franklin Lakes, New Jersey, USA) and the dedicated panels NMIC-402 for all the diagnostic routine samples and NMIC-417 for all the other samples. The antibiotics and the range of antibiotic concentrations present in the two panels are listed in Table S2.

### DNA extraction, sequencing and bioinformatics

Genomic DNA was extracted from all samples using a QIAsymphony instrument (Qiagen, Milano, Italy) and a dedicated kit (QIAsymphony DSP Virus/Pathogen, Qiagen). All the extracts were stored at -80°C. Genomic DNA libraries were prepared using the Nextera XT Library Prep Kit (Illumina, San Diego, USA) following the manufacturer’s protocol. Illumina sequencing was performed at 3 centres: Wellcome Trust Sanger Institute, HiSeq X10, 150 base pair, paired-end reads (bp PE; n = 3418); University of Bath, MiSeq, 250 bp PE (n = 110); MicrobesNG (Birmingham), HiSeq, 200 bp PE (n = 15); resulting in 3543 isolates in total, of which 3483 were found to be of high quality. The Illumina sequence reads were trimmed with Trimmomatic v0.33^30^. SPAdes v3.9.0^31^ was used to generate *de novo* assemblies from the trimmed sequence reads using k-mer sizes of 41, 49, 57, 65, 77, 85 and 93 and with the –cov-cutoff flag set to ‘auto’. The assemblies were annotated using Prokka 1.12^32^. Kleborate v0.4.0^33^ was used to group the isolates into *Klebsiella* species using a mash distance threshold < 0.03 to a representative panel of known species.

### Lineage assignment at the sub-species level

Kleborate was also used to assign Sequence Types (STs) to the isolates, but this was only possible for those species for which multilocus sequence typing (MLST) schemes have been previously established^34^. We therefore carried out intra-species lineage assignments into sequence clusters (SCs) for all species using PopPunk 2.0.2^35^. For each species, the number of components to fit in the mixture model (k) was chosen based on the scatter plot of core and accessory distances. The model was then fit, and the boundary refined using an iterative process of moving the boundary and reassessing the network features. In all cases the core boundary was used to define the clusters. For most species, 2 components provided the best fit, with the exceptions being *K. aerogenes* (k=6), *K. michigenensis, K. quasipneumoniae subsp. quasipneumoniae*, and *K. terrigena* (k=3). For two species there were outlying isolates (SPARK_1532_C1, SPARK_1532_C2; SPARK_1553_C1, SPARK_871_C1) which were removed to fit the model, and then reassigned as query sequences. Plots of the final fits are shown in Figure S1. The SCs defined by PopPunk were named according to their relative abundance within each species, with SC1 being the most abundant, followed by SC2, and so on. For those species where STs could also be called using Kleborate, the SCs defined by PopPunk matched closely with STs (Rand Index > 0.98), although SCs tended to be slightly more inclusive groupings. For ease of reference, we also used a compound identifier that combined SC with the corresponding canonical ST (for example, the most common group in *K. pneumoniae* was designated *K*.*pne*_SC1_ST307).

### Inferring transmission events

To quantify the density of transmission events within and between different sources, we aggregated data from each *Klebsiella* species and identified transmission events by using a threshold-based approach. For each SC, we created a network where all isolates were connected to each other. We then cut this network into ‘putative transmission clusters’ by removing all the links between isolates where the distance was equal to or greater than the threshold. For each of these putative transmission clusters, we recorded every source pair which was connected by a pair of isolates where the distance was below the threshold (including same source pairs), and for each source pair (not isolate pair) which satisfied this we recorded a single transmission event. This approach was conservative because these clusters may in reality have corresponded to multiple transmission events between any two given sources, but it avoided the risk of over-estimating transmission events by counting single events more than once. To obtain transmission frequencies we normalised the count of transmission events by the total number of isolates in each pair of sources. We performed this procedure 4 times, each using a different threshold, to ensure that our results were not overly influenced by the choice of threshold (data not shown).

We used two methods to calculate the distances between isolates; a direct single-nucleotide polymorphism (SNP) distance obtained from mapping to a closely related isolate, and a kmer based core-genome distance estimated by PopPunk. For the mapping based distance, for each SC in the dataset we randomly chose an isolate to use as the reference, and then mapped all isolates from that SC to this isolate using Snippy (https://github.com/tseemann/snippy). We then used SNP-dists (https://github.com/tseemann/snp-dists) to count the number of SNPs between each pair of isolates from that SC. For the PopPunk distance, we used the core genome distance estimation from PopPunk (sketch size 1000000) multiplied by 5000000 (genome size) to obtain an approximate SNP distance between pairs of isolates. We used thresholds of 20 and 50 SNPs, and counted transmission events as described above for each of these thresholds using the two types of distance measure (SNP 20, SNP 50, k-mer 20, k-mer 50).

### Transmission modelling

We used a system-dynamic compartmental model of ordinary differential equations (ODEs) to describe the spread of *Klebsiella* between distinct ‘nodes’, *x* (all sources) within the transmission network with size *N*. Each node *x* can take one of *N* = 24 values, each representing a source. We assume that each node *x* is expressed as the proportion of *Klebsiella* susceptible samples *S*_*x*_ , and the proportion of *Klebsiella* infected samples *I*_*x*_ . Between nodes, β_*xy*_ is the weighted relative transmission from source *x* to sink *y. g*_*x*_ is the recovery/decay/loss of infection from infected *Klebsiella* to susceptible *Klebsiella* and *N* is the total number of nodes (*N* = 24). *b* is the relative scaling for the parameter β _*xy*_.

We adapted the susceptible-infected (SI) model with recovery/decay/loss of infection, expressed as the rate of change over time for *Klebsiella* susceptible and infected samples *S*_*x*_ and *I* _*x*_.

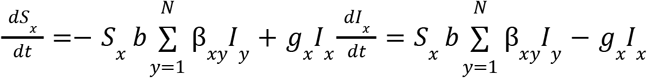

We combined the transmission event and prevalence data to calculate the relative weight of transmission. These weights were used as an indication of the magnitude of transmission between nodes, which informed a general ‘*hierarchy of transmission*’, and accounts for different sample sizes. These weights were calculated as the number of transmission events between nodes, divided by the total samples taken within the ‘from’ node (e.g., if 3 transmission events were observed between human-community carriage and human-hospital carriage from 85 human-community carriage samples then the relative transmission is 3/85 = 0.0353), alongside a 95% credible interval (95%CI = 0.000 – 0.0745). From the prevalence values, we calculated a feasible range of fitting values for each node. This was calculated as +/- 25% of the original prevalence (with an upper limit of 100%). The direction of transmission is unknown therefore we populated the transmission matrix β_*xy*_ with both *x*→*y* and *y*→*x*, with a 50% proportion of each relative weight of transmission.

To capture the underlying uncertainty within parameters, we used Latin-Hypercube sampling to find a credible range of parameters which simultaneously replicated the prevalence data. We varied the following parameters:

- *b*, 1 – 5; scale of transmission.
- *g*, 0.2 – 12 (inverse of 1 month - 5 years range); recovery/decay/loss of infection from infected to susceptible *Klebsiella*.
- Transmission weight from source to sink to populate β _*xy*_, varied between each calculated 95% credible interval (total 71).

For these 1+1+71= 73 parameters, we ran the system of ordinary differential equations for 20 years from 2000 – 2020, with an initial condition of the final 2020 values. We did this for a total of 1000000 parameter sets. Each simulation was either accepted if the equilibria prevalence fell within the range +/-25% for each node simultaneously (if a simulation returned a fit for one node, we assigned a score of 1), or if they failed to satisfy any fitting range, the simulation was rejected.

## Results

### Sequencing, species assignments, and phylogenetic analysis

After quality control, 3,483 high quality read sets and assemblies were retained from the 3,543 libraries; 2,796 from diverse sources recovered using SCAI media, and 687 from ongoing clinical surveillance projects. All except one of these isolates were assigned as *Klebsiella* species by genome sequencing (see below). Summaries of all sequenced strains, including species, lineage assignments and source are given in Table S3. Full details, including the genotypic and phenotypic resistance data, other output from Kleborate, geographical data and phylogenetic trees are available for download via the Microreact project at https://microreact.org/project/KLEBPAVIA. A summary of the main metadata fields used in the Microreact project is provided in Table S4.

Summaries of the species assignments and sources of the 3,483 sequenced isolates are given in Figure 1, as well as phylogenetic trees as described below. We have adopted the three letter species abbreviations used in Figure 1 throughout this paper; a key is provided in the legend. All isolates were assigned as *Klebsiella*, with the exception of a single isolate recovered from the surface of an Automated Teller Machine (ATM) which was assigned as a novel species belonging to the genus *Superficiebacter*, designated *S. maynardsmithii*. This genome, which is described elsewhere^36^, did not contain any notable resistance or virulence features, but was retained as a convenient outgroup. The WGS data confirmed the remaining 3,482 isolates as *Klebsiella*. All of these isolates were initially assigned to one of 7 species by MALDI-TOF, and subsequently assigned to one of 15 species using Kleborate and phylogenetic analysis of the WGS data as described below. The accuracy of the MALDI-TOF assignments varied according to species; 88.4% of the isolates assigned as *K. pneumoniae* by MALDI-TOF were confirmed by WGS, whereas only 30% of the isolates assigned as *K. oxytoca* were confirmed as this species, due to the fact that reference databases are currently unable to distinguish *K. oxytoca* from closely related species^37^ (Table S5).

**Figure 1:**
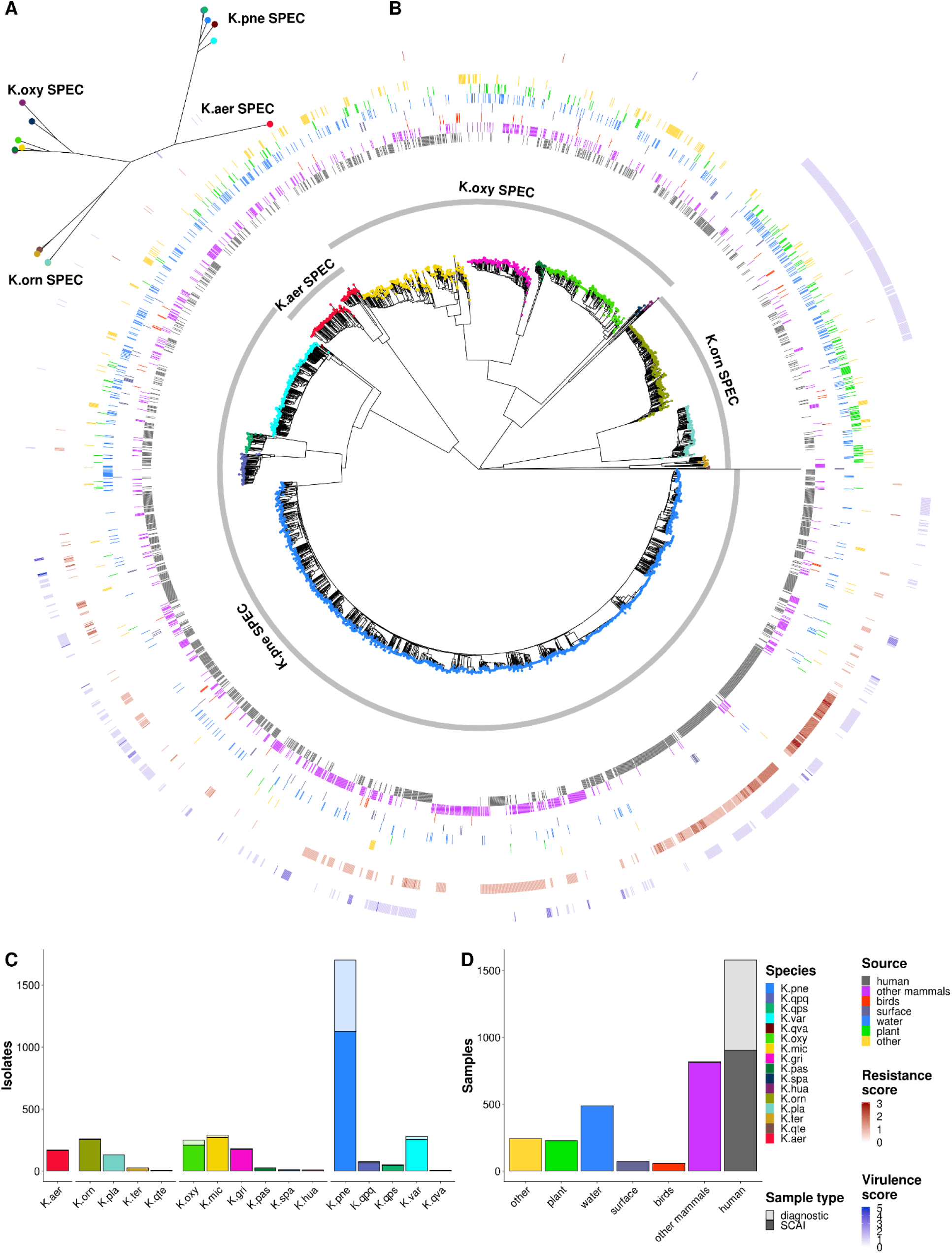
Phylogenetic tree with metadata, and sample and source distributions. **A:** Maximum-likelihood phylogenetic tree constructed from core genes, coloured by species, with the species groups (SPECs) shown. Only one isolate from each species is shown as this tree is intended to show the distances between species. **B:** Neighbour-joining phylogenetic tree constructed from pairwise mash distances between all isolates, coloured by species, with the SPECs shown. The metadata rings show sources (inner rings), and resistance and virulence scores (outer rings). **C:** Bar plot showing the number of sequenced samples from each species. The dark bars show samples from SCAI media, and the transparent ones show diagnostic samples. **D:** Bar plot showing the number of sequenced samples from each high-level source. The dark bars show samples from SCAI media, and the transparent ones show diagnostic samples. The species and species cluster abbreviations are as follows: ***K. pneumoniae Species Complex (K*.*pne*SPEC)**: *K. pneumoniae* (*K. pne*); *K. variicola (K*.*var*); *K. quasipneumoniae subsp. quasipneumoniae* (*K*.*qpq*); *K. quasipnuemoniae subsp. similipneumoniae: K*.*qps; K. quasivariicola: K*.*qva;* ***K*.*oxy*SPEC:** *K. oxytoca; K*.*oxy; K. grimontii K*.*gri; K. michiganensis; K*.*mic; K. huaxensis; K*.*hua; K. pasteurii: K*.*pas; K. spallanzani; K. spa:* ***K*.*orn*SPEC:** *K. planticola; K*.*pla; K. terrigena; K*.*ter; K. quasiterrigena; K. qte; K. ornithicolytica; K*.*orn:* ***K*.*aer*SPEC:** *K. aerogenes; K*.*aer*

We inferred a Neighbour-Joining (NJ) tree of all isolates using Mash distances, and generated a more statistically robust RAxML tree^38^ based on a representative subset of 703 isolates (Figure 1). The Mash-based NJ tree of all isolates is also available to explore and download at the Microreact project given above. The phylogenetic clusters resolved by both the Mash and RAxML trees were entirely consistent with the Kleborate assignments, except for those cases where clusters were not present in the Kleborate database. We assigned the isolates to 15 described *Klebsiella* species, including *K. pasteurii (K*.*pas)* and *K. spallanzanii (K*.*spa*) that were first isolated during the course of this study and are described elsewhere^39^, and eight isolates of the recently described *K. huaxiensis (K*.*hua*) that has previously only been recovered from a urine sample from China^40^ (supplementary note 1). In addition to these 15 recognized species, our data resolve a novel cluster of 6 isolates, to which we have assigned the label *K. quasiterrigena (K*.*qte*), and 2 isolates from hospital carriage that are positioned approximately equidistantly from *K. grimontii (K. gri)* and *K. pas*, to which we have assigned the label ‘NA’ within the Microreact project. Further details on these novel clusters and recently described species are given in supplementary note 1 and their phylogenetic positions are shown in Figures S2 and S3.

*K. pneumoniae* (*K. pne*) is by far the most commonly sampled species, accounting for approximately half of the isolates (n=1705). Previous studies based on WGS data do not support the assignment of the *Raoultella* species as a separate genus^7,26,41^, and this is further supported by our data. Hence in this work we have referred to these species as *Klebsiella*. Both the Mash and RaxML trees revealed that the species resolve four higher-order clusters which we have called species clusters (SPECs; to distinguish from Sequence Cluster (SC)) and named according to the canonical species in each group: *K*.*pne*SPEC, *K*.*oxy*SPEC, *K*.*orn*SPEC, and *K*.*aer*SPEC. The *K*.*pne*SPEC is relatively divergent from the other SPECs, *K*.*aer*SPEC occupies a central position in the phylogeny, and the *K*.*orn*SPEC (the “*Raoultella*” group), forms a sister group to *K*.*oxy*SPEC. *K*.*oxy*SPEC is the most species-rich cluster in our data, but *K*.*orn*SPEC occupies the widest genetic breadth with a single deep division separating *K*.*orn* and *K*.*pla* from *K*.*ter* and *K*.*qte*. Trees for each of these species clusters except *K*.*aer*SPEC are provided in Figures S2-S4.

### Species Clonality and Population Structures

The delineation of isolates into sequence clusters (SCs) by PopPunk (Methods, Figure S1) facilitated a broad comparative analysis of the population structure of each species (Figure 2). Previous genomics datasets have revealed a high degree of lineage diversity within the *K*.*pne*SPEC ^*42,43*^, and our data reveals that this is typical for the four complexes of the genus. In total, we identify 1,168 SCs across all species, of which only 41 (3.5%) are represented by > 10 isolates. Moreover, 50% of all isolates correspond to SCs that are observed no more than 6 times. The most common SC within each species represents between 3-10% of the population (Figure 2a), and most SCs are very rare, hence the SC accumulation curves are not close to saturation (Figure 2b). As shown recently^44^, *K*.*orn* shows particularly high diversity, the 258 isolates of this species resolve into 147 SCs, and the most common SC only accounts for 3% of the isolates. *K*.*pne* has an intermediate level of clonality with respect to the other species; sub-sampling 200 random isolates of this species resolved 95 unique SCs.

**Figure 2:**
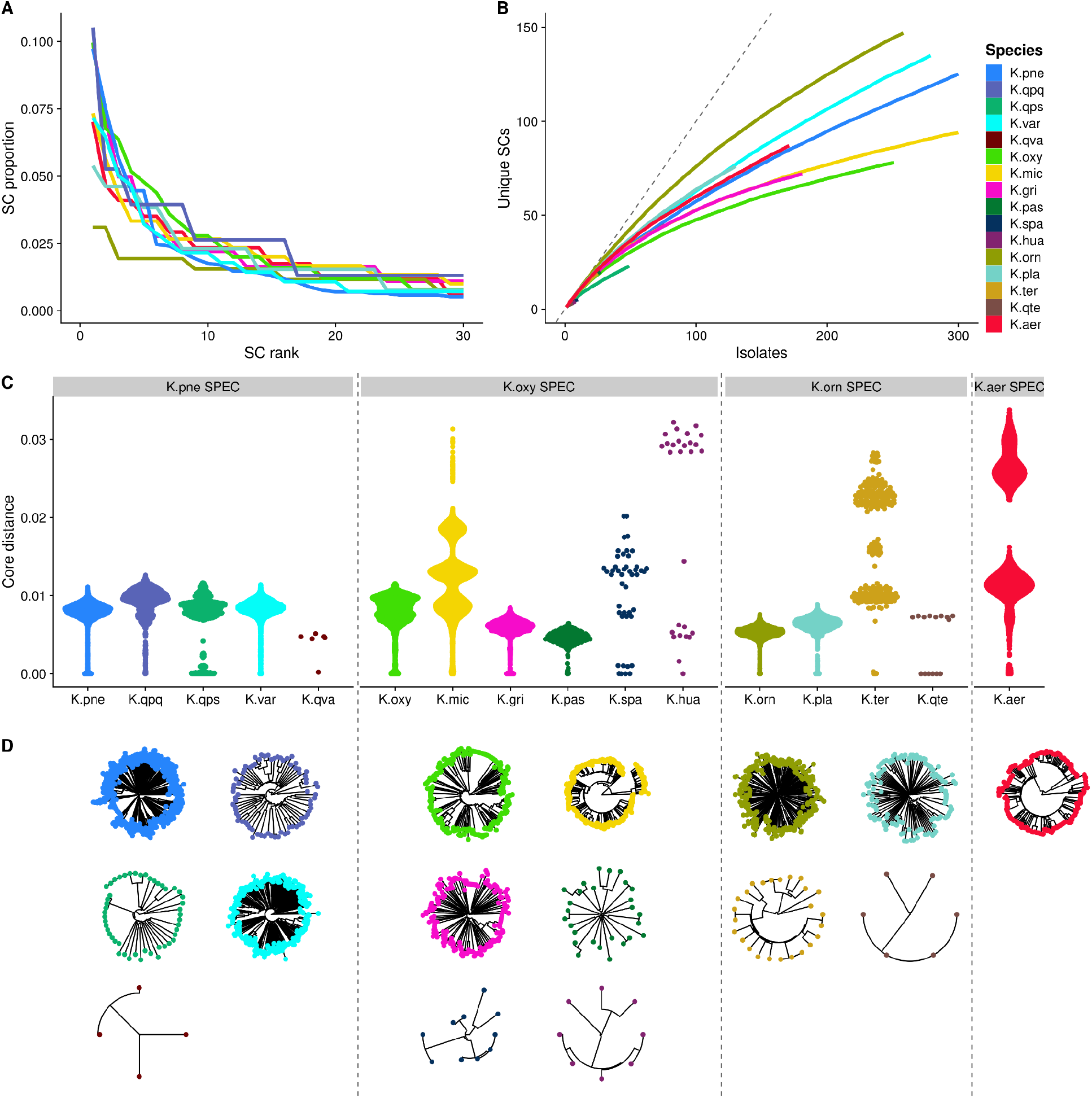
Clonality and Population structure. **A.** The composition of the 8 most common species as determined by SC frequencies. For each species, the isolates were grouped by SC, and the SCs were ranked by their frequencies as a proportion of the dataset (top 30 SCs shown). **B:** The number of unique SCs as isolates are sampled. Accumulation curves were produced by randomising the order of the isolates and counting SCs, and then repeating this 100 times to smooth the curves. The dashed grey line indicates the x=y line. **C:** The distribution of pairwise core genome distances for each species. The distances were estimated using PopPunk, and the points were arranged in the x direction by density to show their distributions. **D:** Neighbour-joining phylogenetic trees for each species. The trees were constructed from pairwise core genome distances estimated by PopPunk.

For the majority of species, pairwise distances are distributed around a modal average of approximately 1% divergence (Figure 2c). A single dominant modal peak reflects a “bush-like” phylogenetic structure evident in the trees, whereby each lineage is approximately equidistant to every other lineage. In some cases (e.g. *K*.*pne, K*.*gri*) a much smaller peak is also evident at a much lower divergence, reflecting expansion of individual SCs. *K*.*mic, K*.*hua, K*.*spa, K*.*ter* and *K*.*aer* also show more diverged modal peaks, with core genome distances up to 3%; this reflects the presence of deep sub-divisions within these species, and this structure is also evident in the individual species trees (Figure 2d).

### The species are non-randomly distributed across different sources

We explored the prevalence and distribution of the 15 recognised *Klebsiella* species and *K*.*qte* across different epidemiological and ecological sources (Figure 1, 3). The analysis presented in Figure 3 was restricted to the 2,795 isolates recovered using the SCAI sampling strategy, and hence excludes the vast majority of disease isolates from hospital patients. The 687 diagnostic isolates are discussed separately in supplementary materials (supplementary note 2, Table S6). A small number (n=24) of the SCAI isolates were also excluded from this analysis on the basis that they were not sampled from one of the major source categories, or that they could not be unambiguously assigned to a *Klebsiella* species.

**Figure 3:**
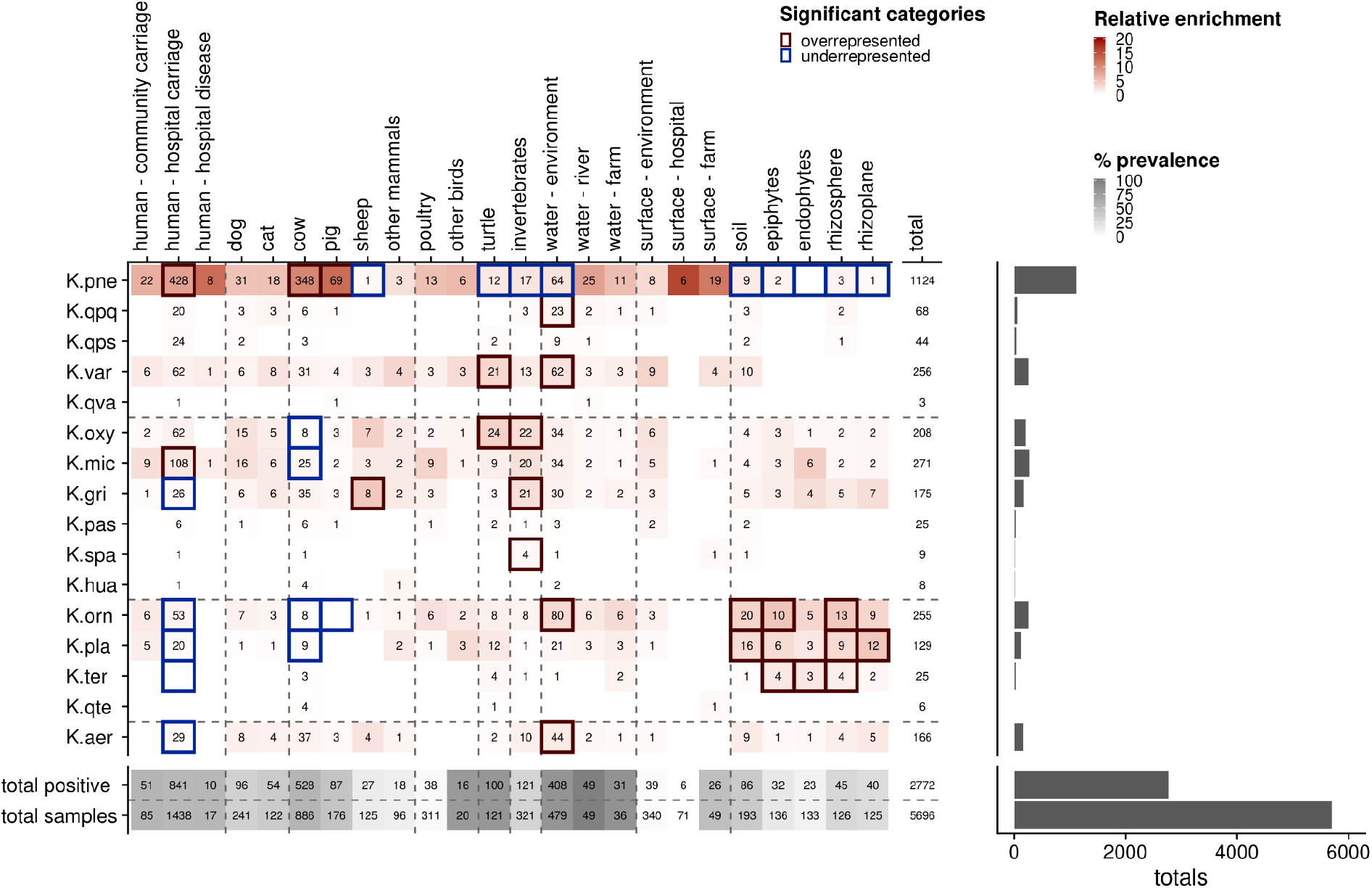
The distribution of species according to source. Only samples from the SCAI dataset (n=2796) are shown, and 24 of these samples were removed either because they were from very poorly sampled sources (21) or could not be confidently assigned to a species (3). The rows represent species delimited according to SPECs, and the columns represent sources delimited according to source categories. The grey shaded rows at the bottom of the table give the total number of positive samples for the corresponding source, and below, the total number of samples for that source. The grey shading reflects the percentage prevalence from each source. The number of positive samples are shown for each species from each source and a blank cell indicates zero positive samples. The red shading shows the relative enrichment of each species from each source, given the overall prevalence from that source, and assuming a null whereby all species would be equally likely to be observed from any given source. The dark red and blue borders show those categories where the number of samples is significantly higher or lower than expected, respectively, as determined by a permutation test. The bar plot to the right shows the number of samples from each species, and the total sampling effort.

Considering all sources, the prevalence of *Klebsiella spp*., calculated as the percentage of samples that were positive for at least one species, was highest for water samples (river, 100%; environmental, 85.2%; and farm 86.1%), as well as turtles (82.6%), most of which live in the river. The source with the next highest prevalence was humans (hospital carriage, 58.5%; community carriage, 62.9%) and livestock (cows, 59.6%; pigs, 49.4%). The prevalence from soil was 44.6% and from plants 26.8%. Whilst a high prevalence was observed from farm surfaces (53.1%), the prevalence from environmental and hospital surfaces was much lower (15.9%). Although the distribution of species across the various sources is clearly non-random (as discussed below), we note that most species can be isolated from most sources; 20 sources harboured at least 7 species, and 11 sources harboured at least 10 species.

We used a permutation test to gauge whether different species are non-randomly distributed between sources (supplementary methods). Significantly non-random distributions are highlighted in Figure 3 (dark red border = significantly overrepresented, blue border = significantly underrepresented). This analysis confirmed that *K*.*pne* is significantly overrepresented in hospital carriage and in livestock (cows and pigs), but is underrepresented in sheep, turtles, invertebrates, environmental water, and soil/plants. In contrast, and as expected, species within the *K*.*orn*SPEC (‘*Raoultella*’) are significantly overrepresented in soil and plants, and underrepresented in hospital carriage. Other species distributions are more surprising. For example, although *K*.*var* is overrepresented in environmental water (in common with *K*.*qpq, K*.*aer* and *K*.*orn*), we do not find any evidence that this species is associated with plants, contrary to its original description^45^.

Interestingly, this analysis also suggests that species from the *K*.*oxy*SPEC tend to be overrepresented in invertebrates, which is consistent with previous reports of a symbiotic relationship between houseflies and *K*.*oxy*^*46*^. Notably, this analysis also points to an overrepresentation of *K*.*mic* in hospital carriage, and we also note a small but significant proportion (18/613; 2.9%) of the diagnostic isolates from hospital disease correspond to this species (supplementary note 2). A caveat with this analysis is that statistical association can result from clonality rather than ecological adaptation. For example, the apparent overrepresentation of *K*.*oxy* in turtles is due to the clonal expansion of a single lineage (SC1) within a population of turtles in a pond at a botanical garden. However, we do not find evidence for clonal expansion of *K*.*mic* within hospital settings, nor for certain *K*.*mic* lineages being more strongly associated with humans than others. Further discussion of the distribution of lineages within species is given below and in Figures S5-S20.

### Distribution of resistance genes

Kleborate^33^ was used to divide all 3482 *Klebsiella* isolates into four resistance scores: 0 = low level resistance, 1 = ESBL positive, 2 = carbapenemase positive, and 3 = carbapenemase plus colistin positive. The distribution of species according to these categories and to each source is shown in Figures 1 and 4. The vast majority of the 3482 isolates (2870/3482, 82.4%) are category 0 (low level resistance). Almost all other isolates are either *K*.*pne* from multiple sources, or isolates of other species from hospital patients, with notable exceptions discussed below. None of the isolates recovered from outside a hospital setting harboured a carbapenemase gene, or showed phenotypic non-susceptibility to carbapenems. This is true for all species, including *K*.*pne*. Only three isolates of species other than *K*.*pne* from outside the hospital setting harboured an ESBL gene (*bla*_SHV-12_ in each case), and one of these (SPARK_2923_C1) is a *K*.*orn* isolate recovered from a fly caught within a hospital, thus pointing to a potential role for fly-mediated AMR transmission within the clinical environment. Excluding *K*.*pne*, there were nine isolates from other species recovered from hospital patients that harboured ESBLs (*bla*_CTX-M-15_, n=4; *bla*_SHV-12_, n=5). Of note are a pair of clonally related isolates (SPARK_1773_C1, SPARK_2031_C1), belonging to clone *K*.*qpq*_SC_11_ST571, that harbour *bla*_CTX-M-15_, plus the virulence factors *ybt, iro* and *rmpA*. These isolates were recovered from urine samples from inpatients at the same hospital in April 2018. This is consistent with hospital transmission of a novel *K*.*qpq* clone exhibiting both resistance and virulence genes.

**Figure 4:**
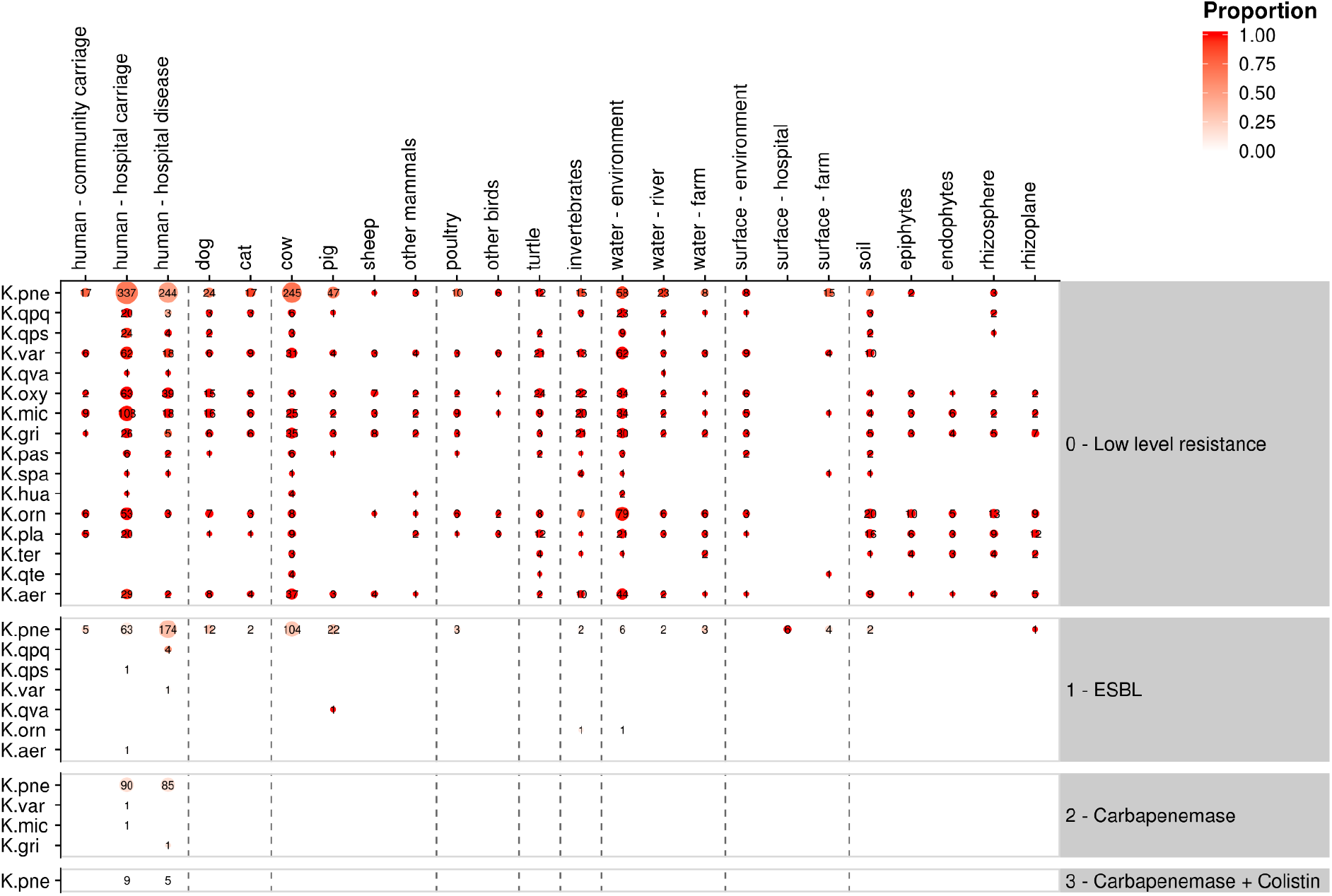
The distribution of resistance genes according to species and source. Resistance genes were identified and grouped into levels 0-3 by Kleborate. The area of the circles is proportional to the number of isolates, and the text shows the number of isolates. The shading shows the proportion of isolates from a given species and source which correspond to a given resistance level.

Excluding *K*.*pne*, only three isolates from other species harboured a carbapenemase gene; these were all isolated from the hospital environment and carried *bla*_VIM-1_. Two of these (*K*.*mic* SPARK_1816_C1 and *K*.*gri* SPARK_1652_C1) present nearly identical genotypic and phenotypic resistance profiles to each other, as well as to five isolates of *K*.*pne*. This resistance profile is characterised by the presence of *bla*_SHV-12_, *bla*_VIM-1_, *mph*(A), and *qnrS* genes, harboured by a class 1 integron (GenBank accession number MN783743) associated with the highly conjugative IncA plasmid pR210-2-VIM^47^. This plasmid is known to be circulating in multiple *Enterobacteriacae* species in Italy^48^, and the re-emergence of VIM-1 in this region is thought to reflect the increased use of ceftazidime-avibactam (CAZ-AZI) to treat with KPC-producing bacteria. Our data provide evidence of inter-clonal and inter-species transfer of this plasmid. Mob-Suite^49^ and Abricate (https://github.com/tseemann/abricate) revealed the presence of a pR210-2-VIM-like plasmid (GenBank accession number CP034084) in unrelated *K*.*pne* clones within a single outpatient (consistent with intra-patient transfer between *K*.*pne* lineages), within different *K*.*pne* clones in two different hospitals (inter-hospital transfer), and also within the isolates of *K*.*gri* and *K*.*mic* from a single hospital (inter-species transfer) (Figure S21). This plasmid is also closely related to the MN78374 VIM plasmid isolated from *E. coli*^*48*^.

Regarding the 1705 isolates of *K*.*pne*, 1105 (64.8%) exhibited a low level of resistance (category 0), 411 (24.1%) carried an ESBL (category 1), 175 (10.3%) carried a carbapenemase gene (category 2) and only 14 (0.8%) carried a carbapenemase gene and colistin resistance (category 3). Two ESBL genes were dominant; *bla*_CTX-M-15_ and variants of *bla*_SHV-27_, which together accounted for 83.5% of all ESBL genes. These were non-randomly distributed between sources; 238/256 (93%) of the *K*.*pne* isolates bearing *bla*_CTX-M-15_ were from humans, the exceptions being from hospital surfaces and companion animals. In contrast, only 51/170 (30%) of the *K*.*pne* isolates bearing *bla*_SHV-27_ variants were from human sources, compared to 87/170 (51%) from cows^50^. Of the 175 *K*.*pne* isolates harbouring carbapenemase genes, all were isolated from the hospital environment and the vast majority (n=161; 92%) carried *bla*_KPC_ and correspond to healthcare associated clones ST258/512 or ST307.

*K*.*pne*_ST307_SC1 was the most abundant clone in the dataset, and was isolated from hospital surfaces and companion animals as well as hospital patients, although none of the ST307 isolates from non-human sources harboured *bla*_KPC_. Eleven *K*.*pne* isolates harboured *bla*_VIM-1_, including those discussed above, and 3 *K*.*pne* isolates harboured *bla*_OXA-48_. Of the 192 isolates with a carbapenemase gene for which phenotypic resistance data is also available, 91% showed phenotypic resistance to ertapenem, 71.7% to imipenem, 77.7% to meropenem. In contrast, the values are 0.8% (27 isolates), 0.18% (6 isolates) and 0.28% (9 isolates) respectively for isolates (from all species) without a carbapenemase gene, and these exceptions are likely to be due to changes in membrane permeability^51^. Consistent with the genotypic data, there is no evidence for any phenotypic resistance to carbapenems outside of the hospital environment.

There were 14 isolates in the highest resistance category; these were all *K*.*pne* isolates from hospitals and all harboured the carbapenemase gene *bla*_KPC_ plus a mutated *mgrB* gene known to confer colistin resistance. All except one of these isolates belong to the common healthcare associated clone ST258/512, with the exception of a single ST307 isolate. Phylogenetic analysis using RAxML of the 95 ST258/512 isolates suggests at least five acquisitions of the *mgrB* chromosomal mutation into this clone (Figure S22). The available phenotypic data confirms resistance to colistin in 12/14 of these isolates. In total, phenotypic resistance to colistin was observed in 46 *K*.*pne* isolates, 41 of which were from humans. Besides the 12 phenotypically resistant isolates harbouring an *mgrB* mutation, Kleborate did not detect a mechanism for colistin resistance in the other cases, including three *K*.*pne* isolates from pigs and a single *K*.*aer* isolate from a goat. This is not unexpected, as many *mcr* variants are not included in the Kleborate database, and colistin resistance can also be conferred through mutations responsible for membrane synthesis^52^. The final non-human colistin resistant isolate was a single *K*.*pne* isolate from a cow that harboured *mcr-1*.

### Distribution of virulence genes

Similar to the genotypic resistance profiles, all isolates were assigned to one of 6 categories based on the presence of the known virulence factors *ybt, iuc, iro* and *clb*, as inferred by Kleborate (Figures 1, 5). In summary, 2749/3482 (78.9%) of all isolates were in the lowest virulence category, a proportion that is slightly lower when only *K*.*pne* isolates are considered (1233/1705; 72.3%). 669/3483 (19.2%) of all isolates correspond to virulence category 1, on the basis that they carry the *ybt* locus (which encodes the siderophore yersiniabactin^53^). The distribution of these isolates varies markedly according to species, corresponding to 410/1706 (24%) of the *K*.*pne* isolates, 249/258 (96.5%) of the *K*.*orn* isolates, 6/279 (2.1%) of the *K*.*var* isolates and 2/171 (1.1%) of the *K*.*aer* isolates. Regarding the strikingly high frequency of *ybt* in *K*.*orn* isolates, we note Kleborate assigns these as an “unknown” type. The *ybt* locus in *K*.*orn* is chromosomally located close to an tRNA-*Asn* site, with no evidence for an associated ICE, and is phylogenetically distinct from the *ybt* locus in *K*.*pne*^*44,54*^. Although this distinct *ybt* locus is essentially core in *K*.*orn*, it is not found in any other species, including other species within *K*.*orn*SPEC, and its function and relevance to virulence in this species remains unclear.

**Figure 5:**
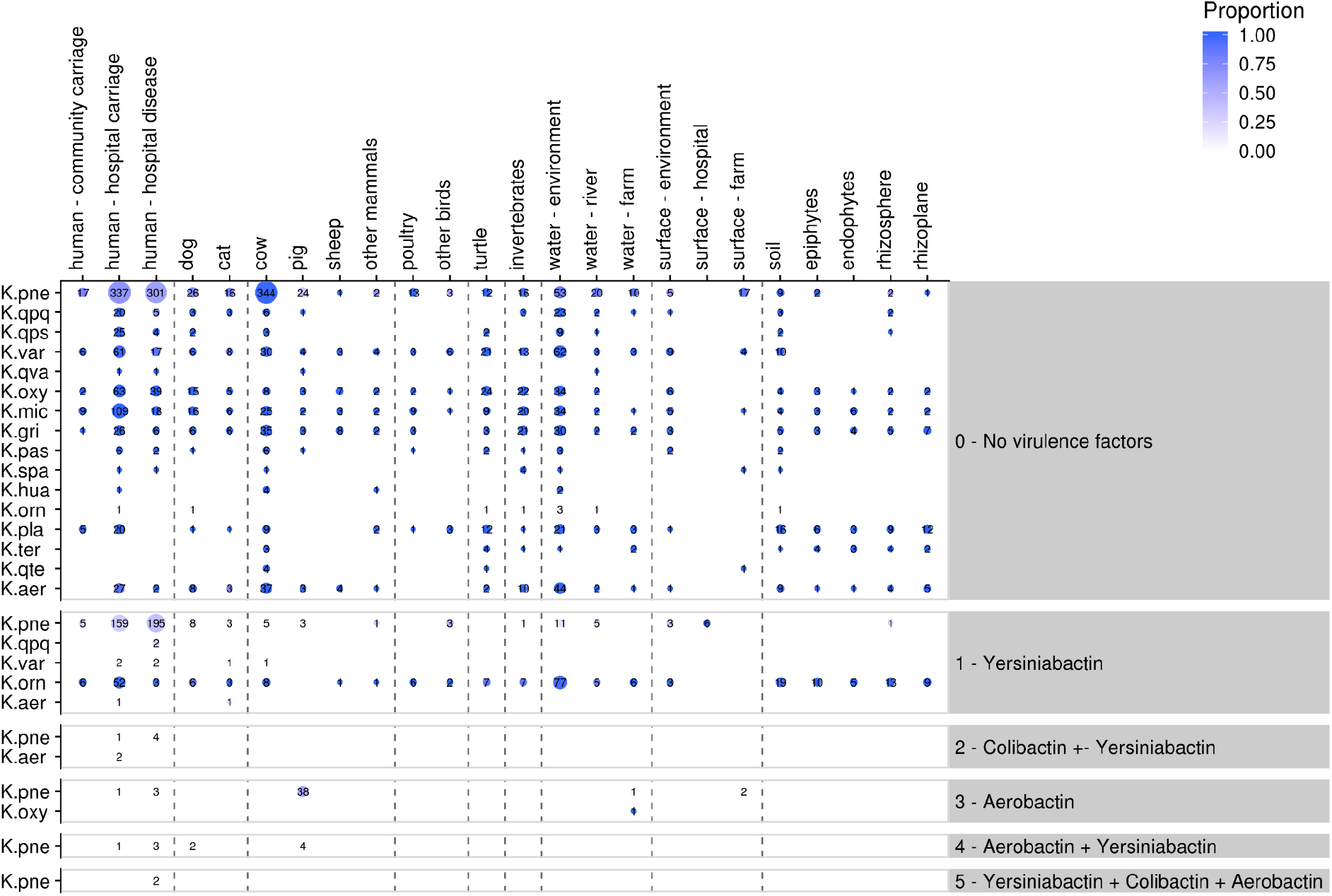
The distribution of virulence genes according to species and source. Virulence genes were identified and grouped into levels 0-5 by Kleborate. The area of the circles is proportional to the number of isolates, and the text shows the number of isolates. The shading shows the proportion of isolates from a given species and source which correspond to a given virulence level.

Whilst only 7 isolates correspond to virulence category 2 (*ybt* plus *clb*), 46 isolates were assigned as virulence category 3 on the basis that they harboured the *iuc* locus that encodes aerobactin. An additional 12 isolates harbouring *iuc* were assigned as category 4 or 5 because they also harboured *ybt*, or *ybt* plus *clb*. All but one of 46 category 3 isolates were *K*.*pne*, with the exception being a single *K*.*oxy* isolate. More surprisingly, 38/46 of the *iuc* harbouring isolates in category 3 were recovered from pigs, and an additional 4 pig isolates from virulence category 4 also harboured an aerobactin. Together, these *iuc* harbouring isolates account for 42/87 (48%) of the pig isolates, and in 40/42 (95%) of these cases the aerobactin locus variant was *iuc*3. A similar association between *iuc3* and pig isolates has recently been described in Germany^55^. The *iuc3* harbouring isolates from pigs represent multiple STs, and were from different farms, hence is unlikely to be due to clonal spread. Moreover, preliminary analysis suggests that it is also not due to the spread of a single *iuc3* harbouring plasmid. Short read contigs harbouring *iuc3* from 48 isolates ranged in length from 55.9 - 430.8 kb and shared between 12.2% and 99.9 % identity. An alignment of six representative contigs from *K. pne* isolates is shown in Figure S23. Three *K*.*pne* isolates and one *K*.*oxy* isolate from the farm environment (water and surfaces) also harbour *iuc3*. Other than the high frequency of the chromosomal *ybt* locus in *K*.*orn*, and of *iuc3* in *K*.*pne* from pigs, the prevalence of virulence genes in livestock and the wider environment was very low.

Twelve isolates were predicted to show a high level of virulence (categories 4 and 5). The 2 category 5 isolates correspond to the hypervirulent lineages *K*.*pne* ST23 and contain all five virulence loci. These isolates were from different hospitals and are not sufficiently closely related to suggest epidemiological linkage. One of these ST23 isolates (SPARK_1158_C1), isolated from the urine of a hospital inpatient, has also acquired the resistance genes *qnrS1* and *bla*_TEM_, and exhibits phenotypic resistance to ciprofloxacin and levofloxacin. We note two additional ST23 isolates from hospitals with a virulence score of 2, and it is known that ICEKp1 is not always present in this clone^56^. Of the 10 *K*.*pne* isolates corresponding to category 4 (the second highest virulence category), eight were from humans and two (representing STs 5 and 25) were from dogs; these isolates harbour *ybt, iuc, iro* and *rmpA*, which points to a risk of transmission of hyper-virulent clones between humans and companion animals.

### The distribution of sublineages below the species level

We examined the distribution across sources of sub-species sequence clusters (SCs), as defined using PopPunk, using the same permutation test as was used to examine species distributions. The distribution of lineages within *K*.*pne* are given in Figure S5 and for other species in Figures S6-S20. As noted above, the species *K*.*pne* is enriched in both humans and cows. However, analysis at the intra-species level reveals that different lineages tend to be associated with either cows or humans, and this is also borne out by phylogenetic analysis (Figure S24). Lineages SC1_ST307, SC2_ST17, SC3_ST512, SC4_ST45, SC11_ST392 are mostly associated with humans, although these vary in the degree to which they are associated with hospital carriage versus hospital disease. For example, considering both SCAI and diagnostic isolates, 66% of the SC1_ST307 (the most common lineage in the dataset over all species, n=166) are associated with disease and 28% with hospital carriage. The equivalent figures for *K*.*pne*_SC2_ST17 (the second most common lineage in our dataset, n=128) are 20% and 57% respectively. A different set of SCs are associated with cows (e.g. SC5_ST661, SC9_ST3068, SC10_ST2703, SC17_ST3345). Some intermingling does occur, particularly in SC5_ST661 which contains clonal expansions of both bovine and human isolates. This lineage has previously been observed from both human and bovine sources^19^, and may represent a more generalist clone that is capable of intraspecific transmission both in cows and humans, but also to occasionally transmit between these two host species.

Fewer statistically significant sequence cluster enrichments are apparent for other species, predominantly owing to smaller sample sizes. However, a number of observations are notable. As discussed, *K*.*mic* appears to be enriched within hospital carriage (supplementary note 2, Table S6), but sequence cluster analysis confirms that this is not due to the expansion of a single SC. Twenty-five of the 30 most common SCs of this species are present in hospital carriage samples, but no single SC is significantly more commonly associated with hospital carriage relative to the others (Figure S11). In contrast, the association of *K*.*gri* with invertebrates appears to be largely driven by the strong enrichment of *K*.*gri* SC1 from this source (Figure S12). This is unlikely to reflect stochastic clonal expansion or sampling bias, as *K*.*gri* SC1 is associated with different invertebrate hosts (a cockroach, fly, wasp and an unspecified ‘bug’) sampled in different locations. This clone, which has no notable resistance or virulence attributes, was also recovered from a cockroach caught in a hospital environment, and an isolate very closely related to this clone was recovered from an outpatient of the same hospital (Figure S25). Similar to the example above of a *K*.*orn* isolate from a hospital caught fly harbouring an ESBL gene, this points to the possibility of invertebrates playing a role in transmission to humans in clinical settings.

### Transmission modelling

In order to quantify and compare transmission events between different settings we used a threshold-based approach to infer transmission, as described in the methods. It is clear from the resulting transmission matrix (Figure 6A) that the vast majority of transmission occurs within a single source and, most importantly, that the vast majority of acquisition by humans originates from other humans rather than from animals or the environment. In particular, our analysis further reinforces the view that transmission of *K*.*pne*, and other species, between cows and humans (which are the two most deeply sampled sources) is limited. Despite this, we note that sporadic transmission events occur relatively commonly between humans and companion animals, and between humans and water sources.

**Figure 6:**
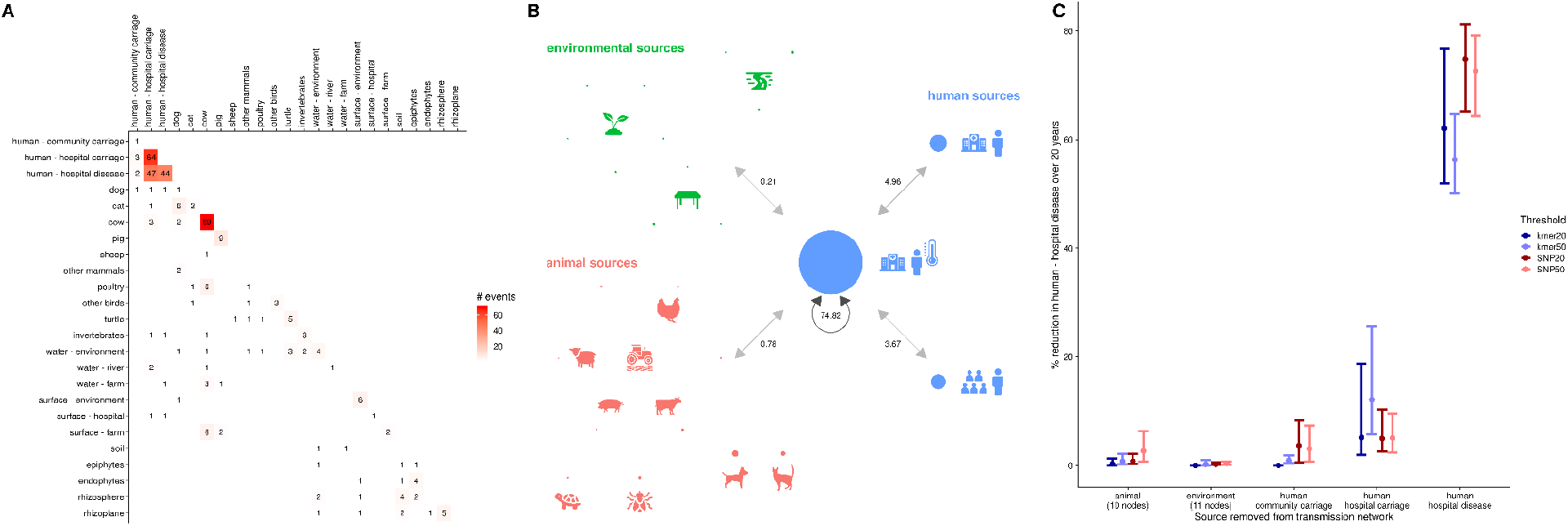
Transmission network modelling analysis. **A.** The number of transmission events between each pair of sources, as determined by a threshold of 20 SNPs. The shading is proportional to the number of events and does not account for the number of samples from each source. **B**. Transmission network showing the impact of removing each node in the network on the prevalence of human hospital disease (the central node). The network shown was produced using a threshold of 20 SNPs. The circles represent nodes, and the area of each circle is proportional to the percentage reduction in hospital disease when that node is removed. The nodes are coloured by the three major groups of sources (human, animal, environment), and the icons give a visual description of how the sources are represented by nodes. The grey arrows show the impact of groups of sources, for example the combined impact of removing all animal sources is a 0.78% reduction in human hospital disease. **C**. Summary of the results from 4 different model runs, each using a different distance threshold to determine transmission events. The points and bars represent the median ± 95% credible interval. The distance estimation methods and thresholds used were SNP distances from mapping to a close reference genome (20 SNPs - dark red, 50 SNPs - light red), and a core genome kmer derived distance from PopPunk (approximately 20 SNPs -dark blue, approximately 50 SNPs, light blue).

To provide a more quantitative assessment of the rates of transmission between different sources we created a system-dynamic compartmental model (see methods, supplementary note 3, figures S26-S29 and Tables S7-S9 for details). We used this model to perform an intervention analysis whereby we removed each transmission link between pairs of sources (including within-source transmission), one at a time, and explored the impact this intervention would have (after 20 years) on the prevalence of *Klebsiella* in the human - hospital disease source (Figure 6B). This analysis showed that the most important source of human infections in hospitals is transmission from other patients with suspected or confirmed *Klebsiella* infection; removal of this transmission edge resulted in a median decrease in prevalence of 74.8% in the human - hospital disease node. The next most important sources were human carriage samples from either the hospital or community; these caused a reduction of 4.96% and 3.67%, respectively, in the prevalence of the human - hospital disease node when their transmission events were removed. We refitted the model and repeated the intervention analysis separately on four sets of transmission events, each calculated using a different threshold (SNP 20, SNP 50, k-mer 20, k-mer 50), to check that our results were not biased by the choice of threshold (Figure 6C). The results from each model run were very similar across the range of thresholds and distance estimation methods, suggesting that our results are robust to these choices.

The suggestion that carriage is a relatively minor factor compared to infected patients in maintaining a high prevalence of *Klebsiella* infection in the hospital is inconsistent with reports demonstrating that gut colonisation is a major risk factor for infection in ICU patients ^57,58^. It is possible that our result in part reflects the high frequency of nosocomial clones in Northern Italy (e.g. K.pne ST307, ST258/512 harbour *bla*_KPC_) that are known to transmit from patient to patient, and from hospital to hospital^27^. Alternatively, this may also in part be a consequence of inflated estimates for the prevalence of hospital disease due to ascertainment bias favouring the recovery of isolates from patients with disease (Berkman’s paradox), hence we urge some caution in downplaying the role of hospital carriage.

Nevertheless, whilst our model suggested that spillover events from animals and the environment do occasionally occur, they contribute very little to the prevalence of *Klebsiella*-associated disease in the hospitals (0.78% and 0.21% reductions for animal and environmental sources, respectively).

## Discussion

The one-health framework is integral to many AMR research programs which aim to mitigate the risks posed by non-clinical reservoirs of AMR through careful surveillance and stewardship. These risks need to be assessed on multiple levels, the simplest being that posed by sporadic transmission events, for example from livestock to farmers, from companion animals to their owners, or from certain recreational activities such as ‘wild swimming’. If these events are epidemiologically distinct, are not dominated by specific lineages associated with heightened virulence or resistance properties, and onward human-to-human transmission is unlikely, then the risk posed to public health by each individual event is likely to be low. There are multiple examples in our data of such sporadic transmission events between humans and animals, and these underpin our transmission modelling. The isolation of the widespread healthcare associated clone ST307 from companion animals, the recovery of highly virulent strains from dogs, and the putative transmission of strains and plasmids between humans and invertebrates in the hospital environment are pertinent examples. Combined with multiple examples from the literature ^59–63^, our data therefore underlines the importance of basic hygiene measures, particularly with respect to contact with animals.

A different, and more complex, question relates to the emergence of high-risk lineages that combine heightened virulence or resistance attributes with the ability to move between, and spread within, different settings. A full understanding of the emergence of such clones requires a consideration of likely anthropogenic drivers, such as inappropriate use of antibiotics or other environmental stresses, but also a broad ecological context. For example, transmission of migrant or emergent lineages within a rugged selective landscape that is characterised by local adaptation and specialist lineages will tend to be relatively restricted. Our data generally reveal low levels of resistance and virulence genes outside of clinical settings, and within species other than *K*.*pne*. Although this may in part reflect biases in the database used by Kleborate towards well-characterised genes known to be common in *K*.*pne*, this observation is also consistent with the view that the emergence and subsequent spread of highly virulent and/or resistant lineages within the environment is a rare event. In contrast, our data provide evidence for the emergence of novel and potentially high-risk lineages within the hospital setting, one example being *K*.*qpq* ST571, that harbours both resistance (*bla*_CTX-M-15_) and virulence (*ybt, iro* and *rmpA*) genes. A second example is the interspecies transfer of the plasmid pR210-2-VIM-like carrying *bla*_VIM-1_ from *K*.*pne* to *K*.*mic* and *K*.*gri* isolates, again within the hospital environment, as well as the intra-patient transfer of this plasmid between different *K. pne* lineages. Our data also reveal a surprisingly high rate of *K*.*mic* within hospital carriage; although in this case there is no evidence for the hospital spread of high-risk *K*.*mic* lineages, the high frequency of this species in hospital carriage, combined with the recovery of a *K*.*mic* strain harbouring the *bla*_VIM-1_ , as well as previous reports of strains harbouring carbapenemase genes^64^ urges heightened clinical awareness of this species.

Our data and analyses thus broadly challenge the view that AMR can ‘flow’ unimpeded between different settings, and we argue that local adaptation plays a role in mitigating transmission. We note that species and sequence clusters within species are non-randomly distributed, with the clearest example being the distinct sets of *K*.*pne* sequence clusters of human and bovine origin, which is consistent with previous studies ^19^. Moreover, modelling of transmission events reveals that transmission is much more common within, than between settings, and that the vast majority of cases of acquisition of *Klebsiella* by humans is from other humans, which is also consistent with previous studies^65^. Nevertheless, several lines of evidence point to companion animals as posing a relatively high transmission risk, as previously reported for *K. pne*^*66*^.

The ecological and phylogenetic distribution of virulence and resistance genes also points to barriers to transmission between humans (and the clinical environment in particular), and animals and the environment. High levels of virulence and / or resistance tend to be rare in species other than in *K. pne*, and outside of the hospital environment. The complete absence of genotypic or phenotypic evidence for carbapenem non-susceptibility outside of health-care settings is particularly striking. Interesting exceptions in terms of virulence include the high frequency of aerobactin in *K*.*pne* isolates from pigs, and the acquisition of a variant yersiniabactin (*ybt*) by *K*.*orn* as a core locus. Although further work is required to explore why these genes are selectively maintained, and their potential risk to public health, the high frequency of these genes in their respective hosts or species suggests an adaptive function.

In conclusion, here we describe whole genome sequence data incorporating multiple species of the *Klebsiella* genus from diverse sources. Our analysis suggests that high levels of resistance and virulence are largely restricted to *K*.*pne*, and that local adaptations may limit the spread of resistant or virulent clones across humans, animals and the environment. Our findings broadly corroborate recent research indicating hospitals as the hubs of *K*.*pne* resistance dissemination in Europe^5^ and justify a continued focus on breaking the transmission chains throughout the health-care network. We add the caveat that ascertainment bias combined with the high frequency of hospital adapted *K. pne* ST307 and ST258/512 in this region (that commonly harbour *bla*_KPC_) will inflate the significance of nosocomial transmission, and infection originating from diverse carriage strains is also known to be common^57^. Moreover, our analysis suggests that novel emergent lineages of heightened virulence and/or resistance are most likely to emerge within hospital settings rather than in the environment or animals, although this possibility cannot be discounted. Our analyses also point to higher rates of transmission within specific animal hosts (eg cows, pigs and turtles) and between plants, although there are some clear routes of transmission between animals and the environment (e.g. between river water and turtles, and livestock and farm surfaces; Figure S28).

We contend that the one-health perspective remains pertinent for restricting sporadic transmission events, and that transmission dynamics will vary according to the region and the pathogen under study. For example, contact between humans and animals may be much more common in many low-resource regions^67^. Finally, we acknowledge limitations with our sample with respect to the role of wastewater and food-borne transmission, and that may play an important role in the transmission cycle between humans, animals and the environment.

## Acknowledgments

This work was funded by the SpARK project, awarded to EF, “The rates and routes of transmission of multidrug resistant *Klebsiella* clones and genes into the clinic from environmental sources,” which has received funding under the 2016 JPI-AMR call “Transmission Dynamics” (MRC reference MR/R00241X/1); and by the French Government’s Investissement d’Avenir program Laboratoire d’Excellence “Integrative Biology of Emerging Infectious Diseases” (ANR-10-LABX-62-IBEID). JC and HT were funded by the ERC grant no. 742158. JC and TK were funded by the Norwegian Research Council grant no. 271162. The use of MRC-CLIMB^68^ was critical for the computational aspects of this work. We are grateful to Ruth Zadocks and Alan McNally for advice during the course of the project.

## Author contributions

HT carried out extensive bioinformatics and statistical analysis, helped write the paper and to design the study. RB developed and implemented the modelling with input from LM and RR. TK, MG, NC, VP, JSLF, CR, SD, FC helped with data analysis and manuscript preparation. CM, MC, CF carried out the sampling, microbiology and susceptibility testing. PM helped with the clinical sampling logistics. SB, DS, JC and EF designed the study, contributed to the analysis and prepared the paper.

## Data availability

Short-read data available under accession numbers ERR3412430 to ERR3412448;ERR3440341 to ERR3440427;ERR3448863 to ERR3449598;ERR3469775 to ERR3469909;ERR3479903 to ERR3480717;ERR3844616 to ERR3844777;ERR3904469 to ERR3904709;ERR3931787 to ERR3932313;ERR3967745 to ERR3967936;ERR4022833 to ERR4023150;ERR4139181 to ERR4139191;ERR4374646 to ERR4374837

Metadata and the tree file can be downloaded from the microreact project at https://microreact.org/project/KLEBPAVIA.

## Code availability

Code for the transmission analysis and permutation tests are available on request.

