## Supplementary figures for "One Health or Three? Transmission modelling of *Klebsiella* isolates reveals ecological barriers to transmission between humans, animals and the environment"

Figure S1

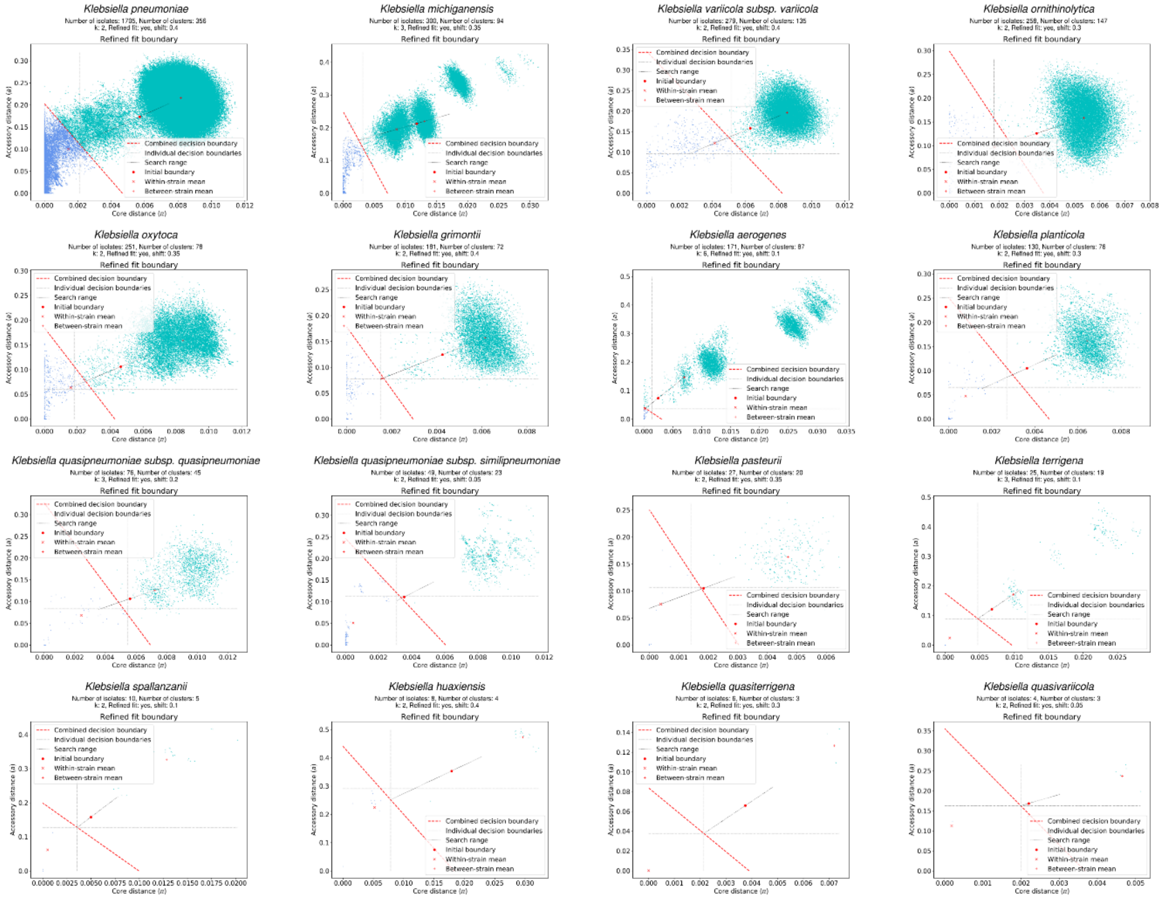


Definition of sequence clusters (SCs) using PopPunk[31]. For each species, the number of components to fit in the mixture model (k) was chosen based on the scatter plot of core and accessory distances. The model was then fit, and the boundary refined using an iterative process of moving the boundary and reassessing the network features. The core boundary was used to define the clusters. For most species, two components provided the best fit. In those species where MLST schemes were available we used the Rand Index to show very high concordance between SCs and STs (RI>0.98).

Figure S2

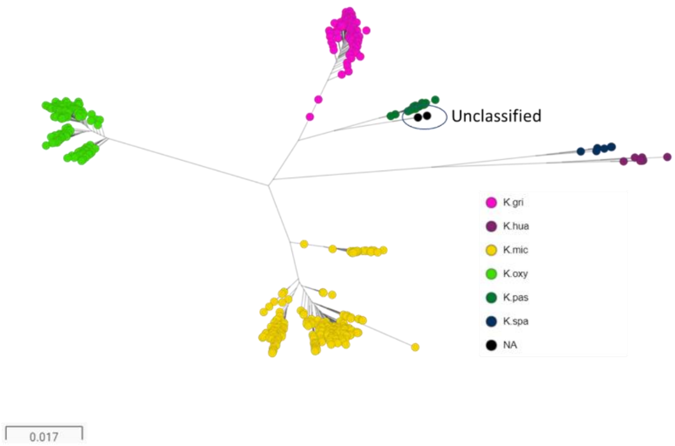


Phylogenetic tree for *K. oxytoca* and related species within the species cluster *K.oxy*SPEC. The tree is generated using MASH distances.

Figure S3

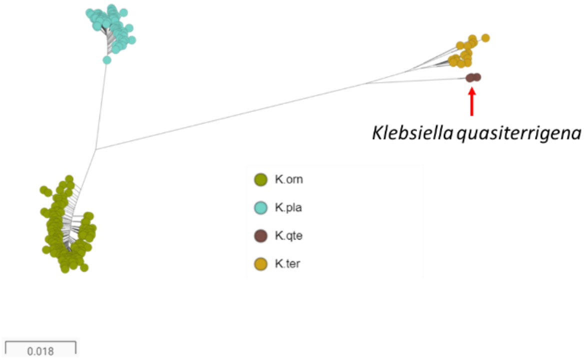


Phylogenetic tree for *K. ornithinolytica* and related species within the species cluster *K.orn*SPEC. The tree is generated using MASH distances.

Figure S4

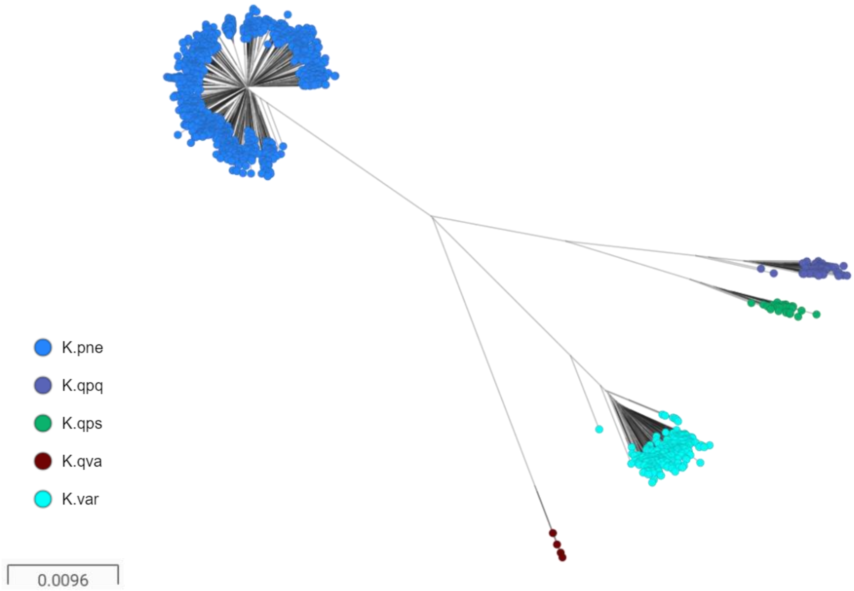


Phylogenetic tree for *K. pneumoniae* and related species within the species cluster *K.pne*SPEC. The tree is generated using MASH distances.

Fig S5 Distribution of *K. pnuemoniae* lineages across different sources

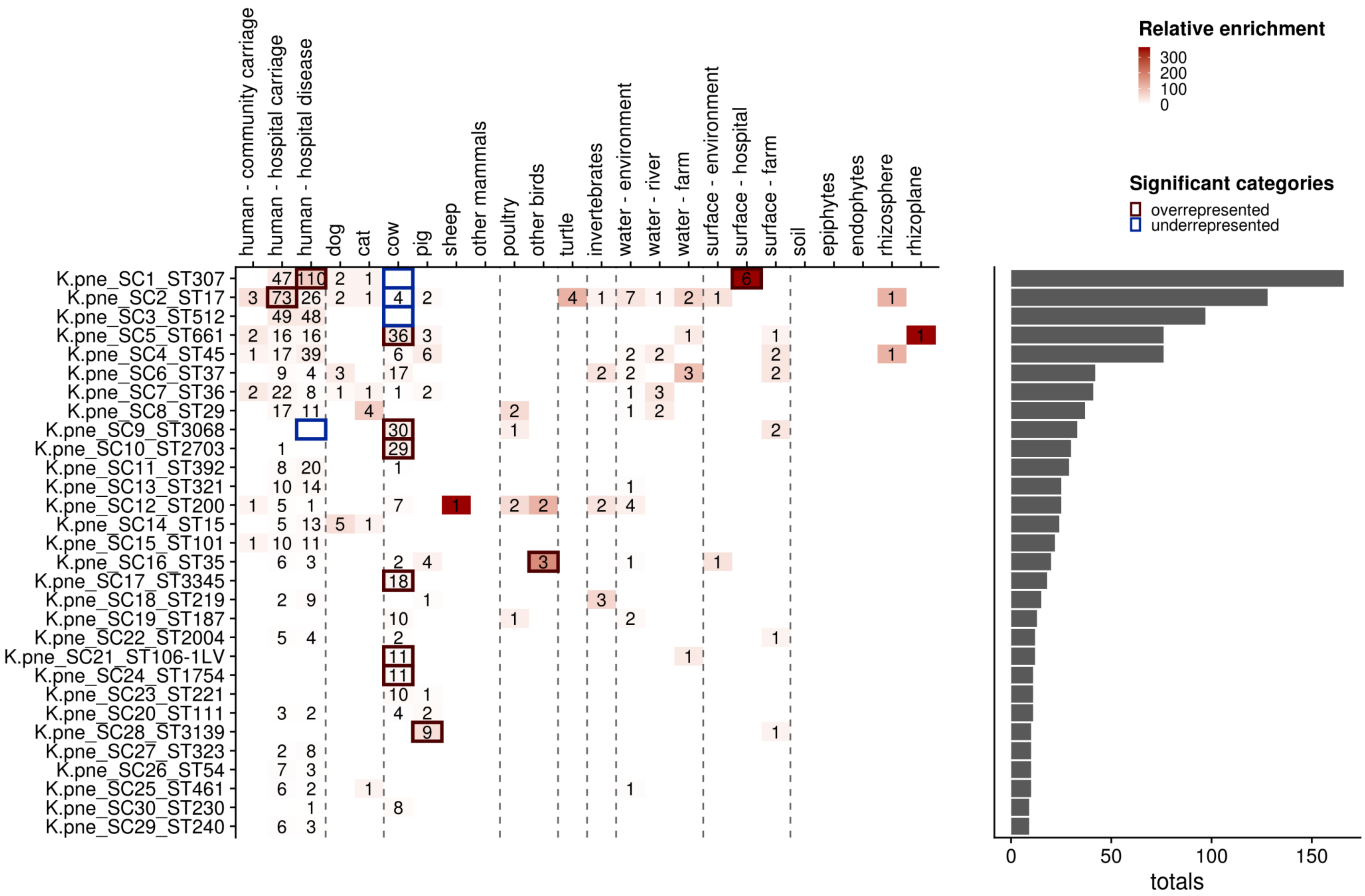

Fig S6 Distribution of *K. quasipneumoniae subsp. quasipneumoniae* lineages across different sources

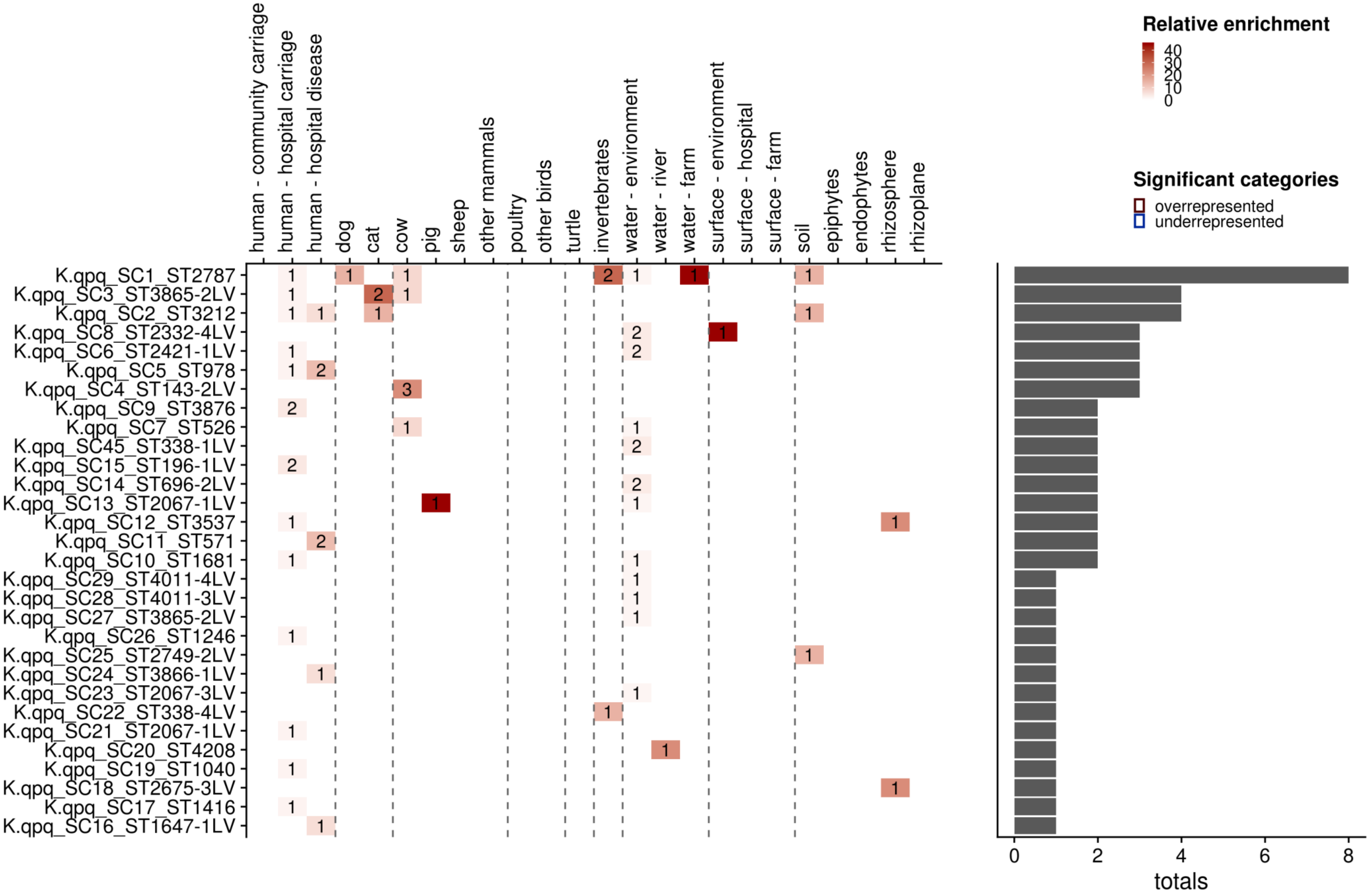

Fig S7 Distribution of *K. quasipnuemoniae subsp. similipnuemoniae* lineages across different sources

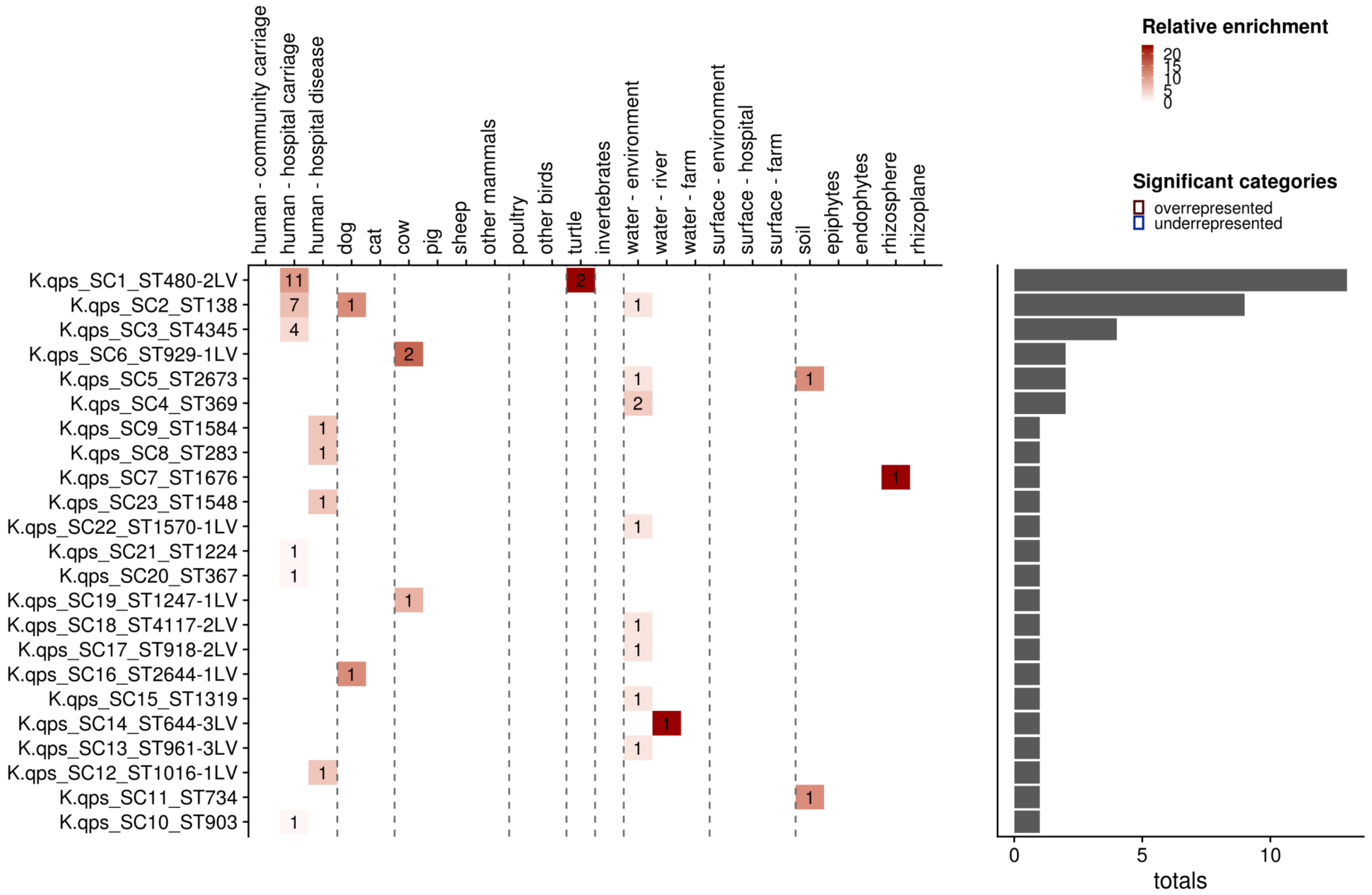

Fig S8 Distribution of *K. variicola* lineages across different sources

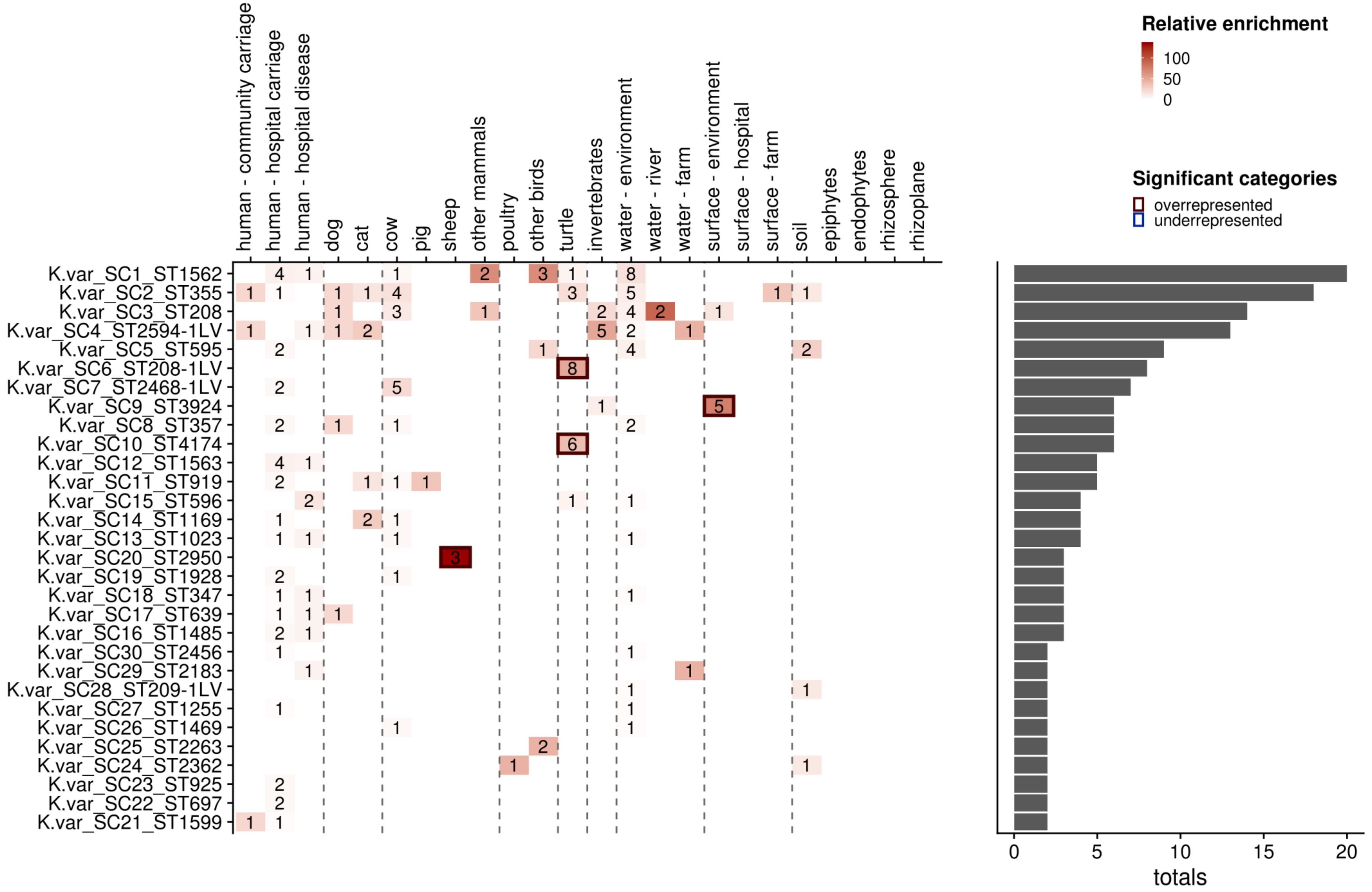

Fig S9 Distribution of *K. quasivariicola* lineages across different sources

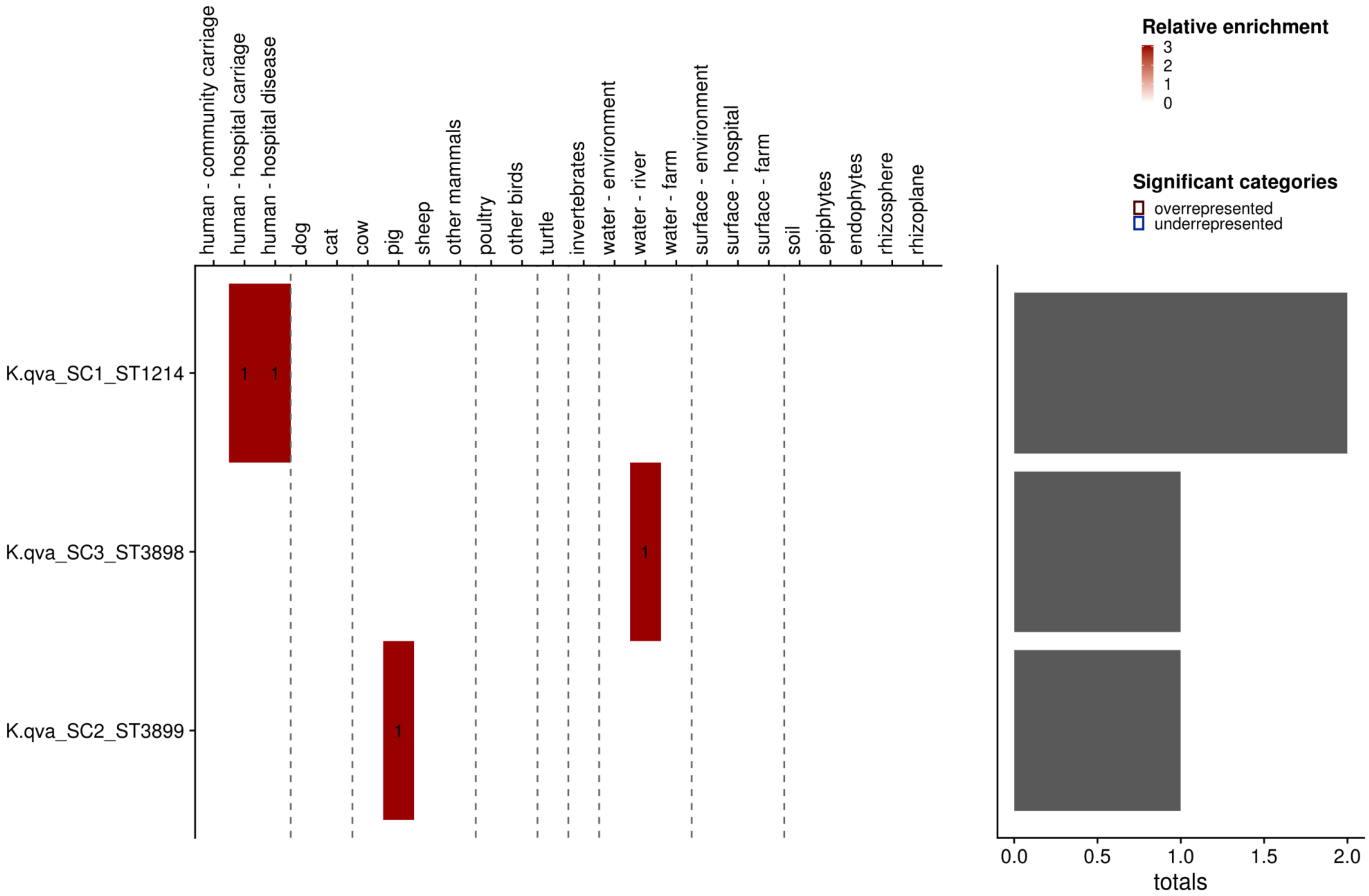

Fig S10 Distribution of *K. oxytoca* lineages across different sources

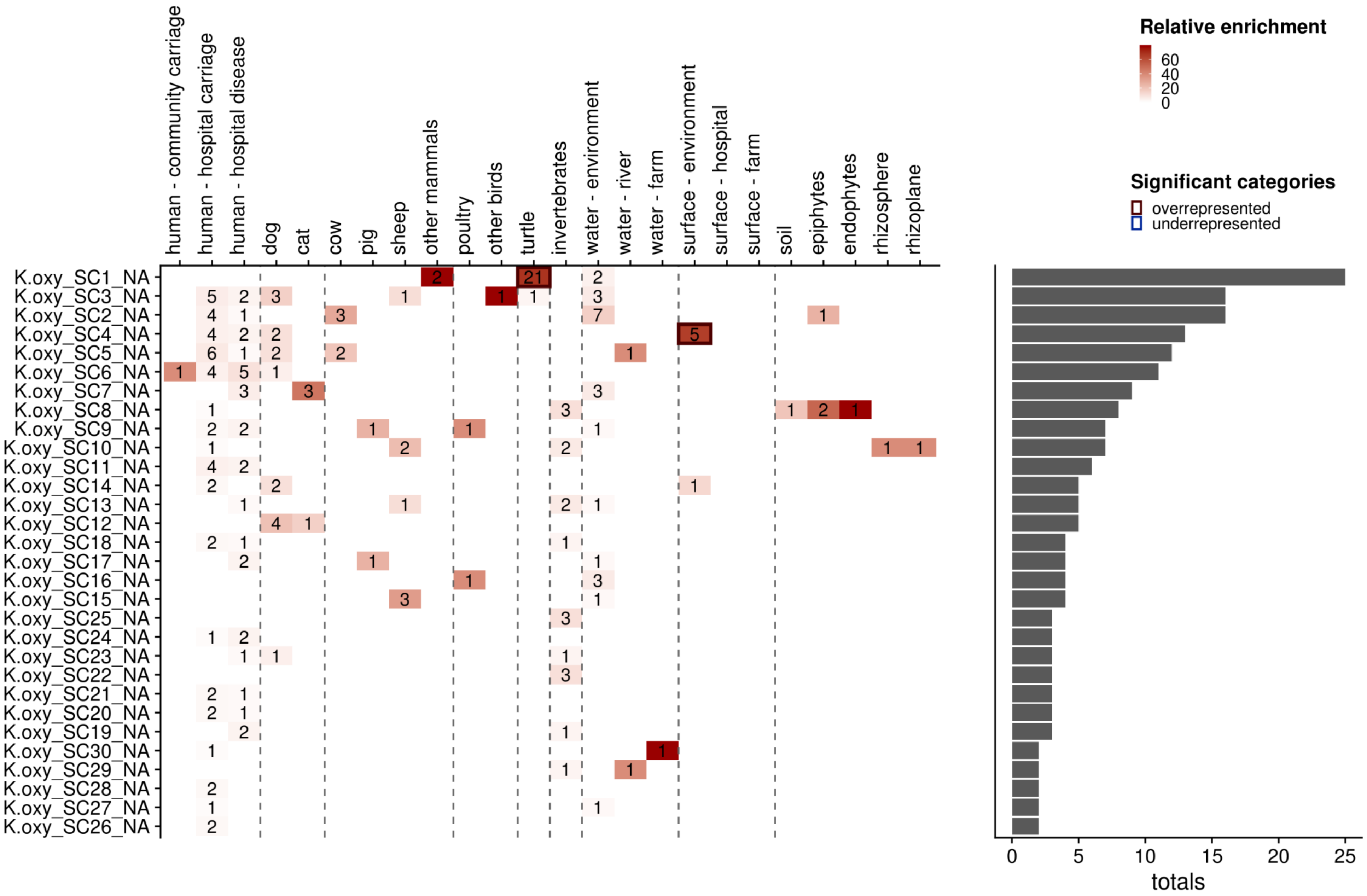

Fig S11 Distribution of *K. michagenensis* lineages across different sources

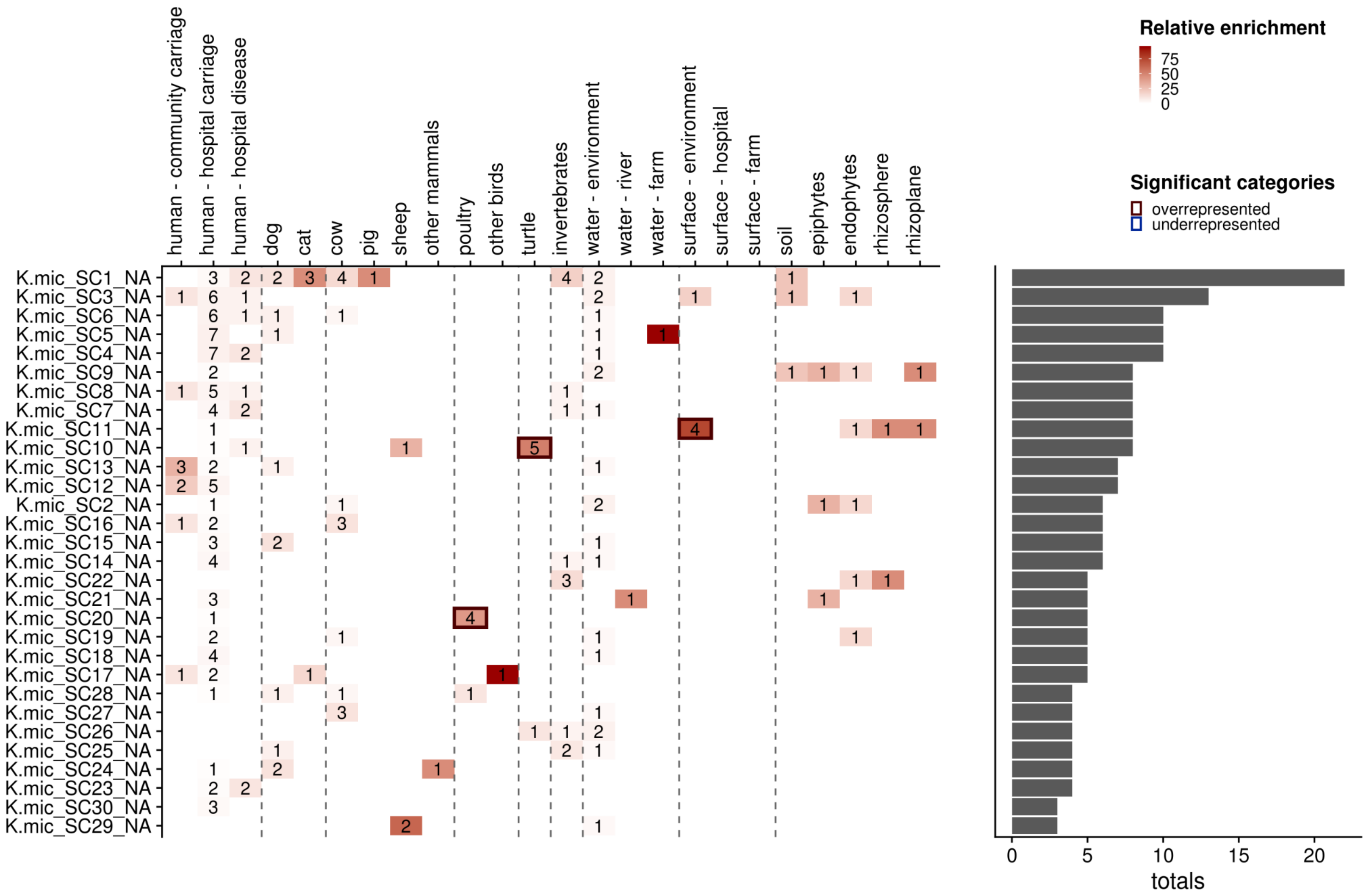

Fig S12 Distribution of *K. grimonti* lineages across different sources

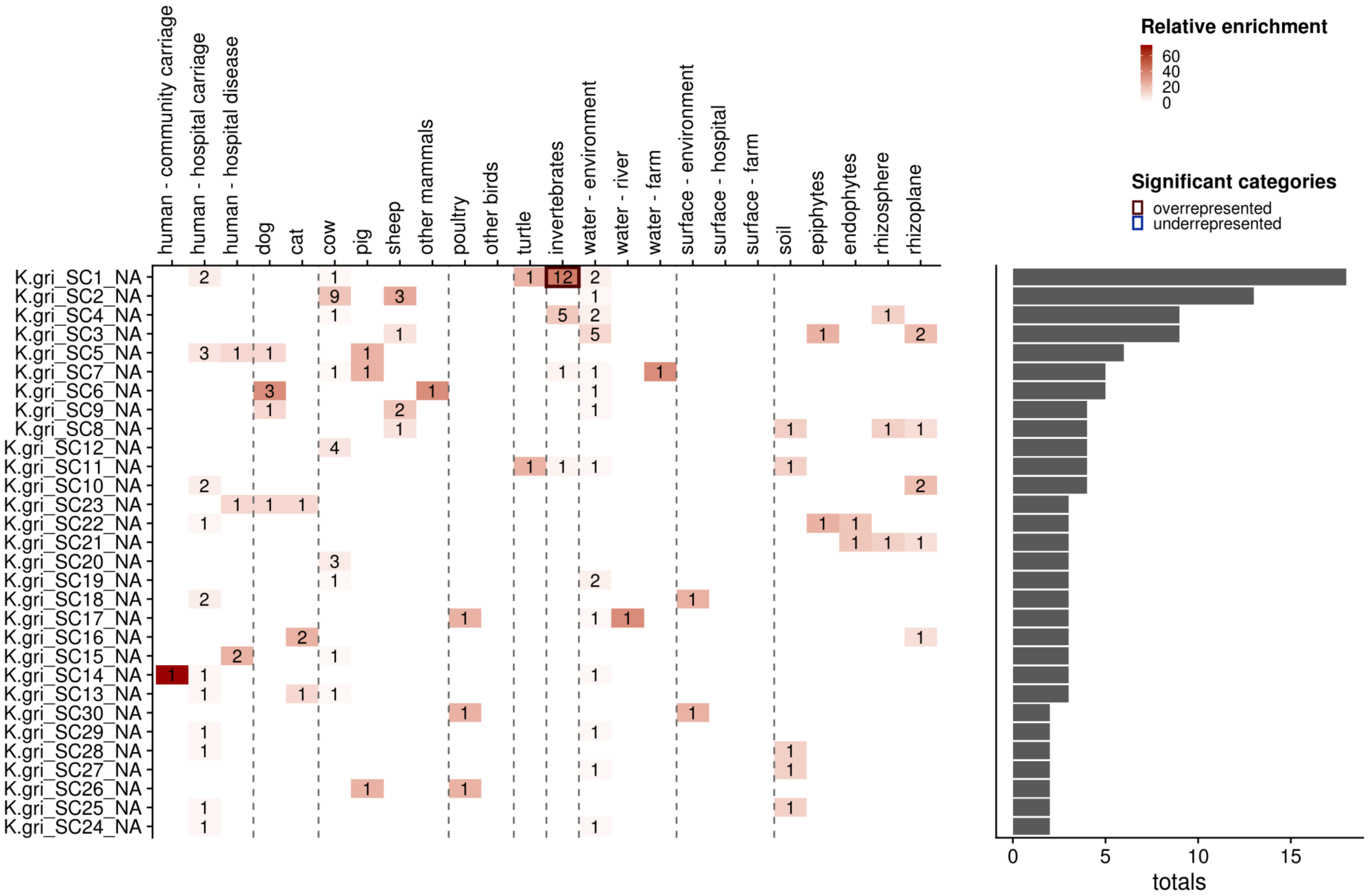

Fig S13 Distribution of *K. pasteuri* lineages across different sources

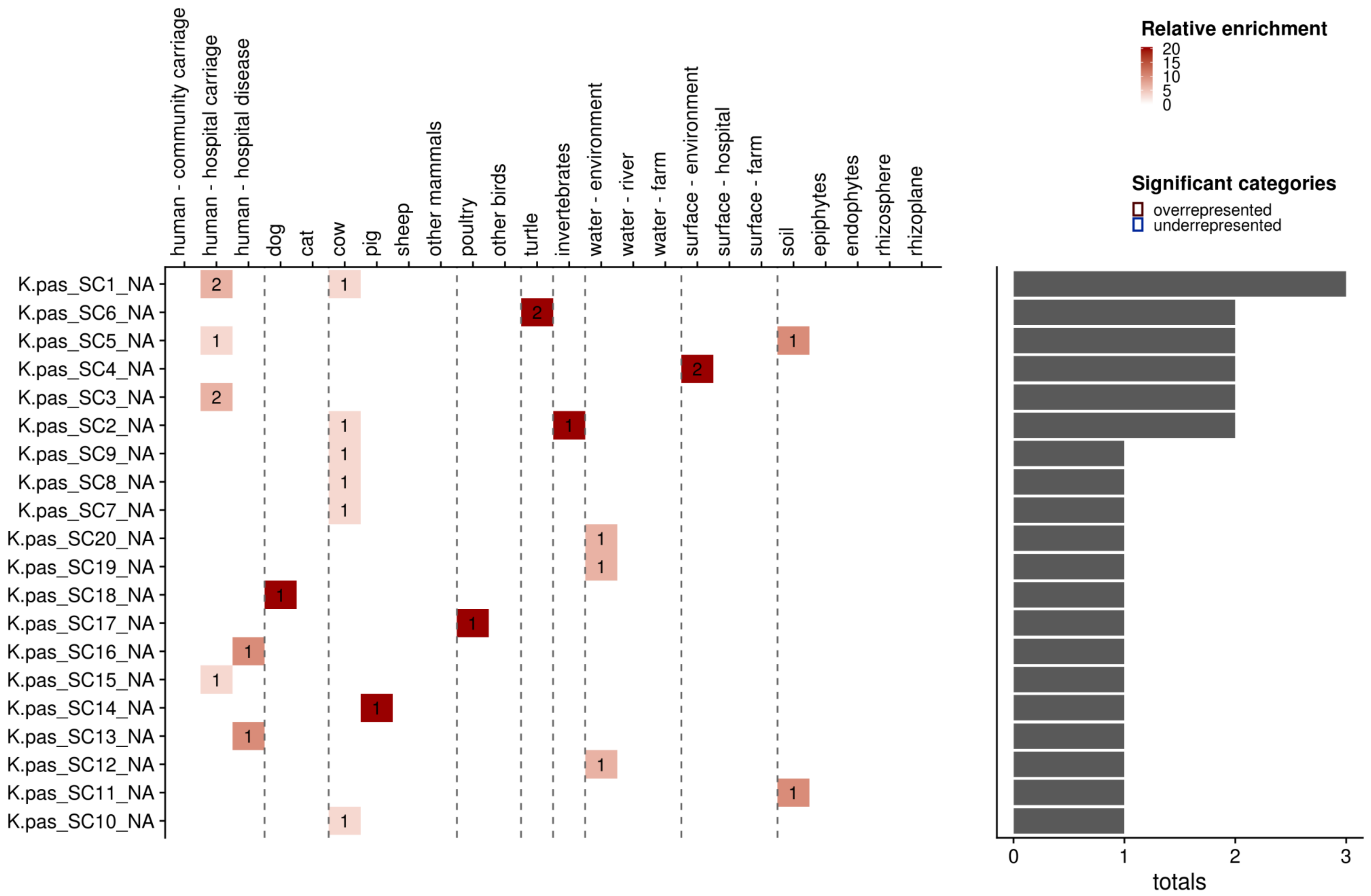

Fig S14 Distribution of *K. spallanzani* lineages across different sources

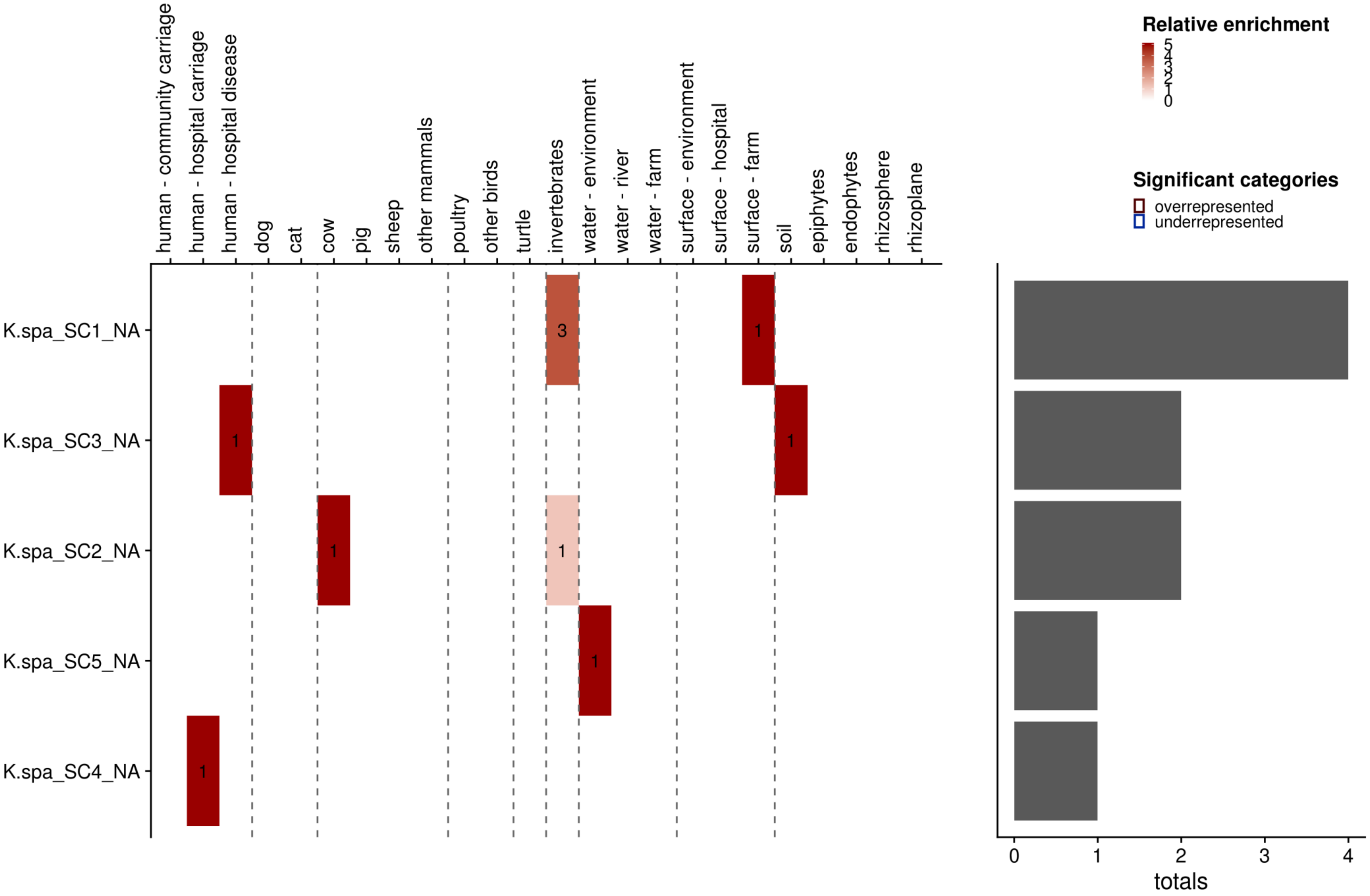

Fig S15 Distribution of *K. huaxensis* lineages across different sources

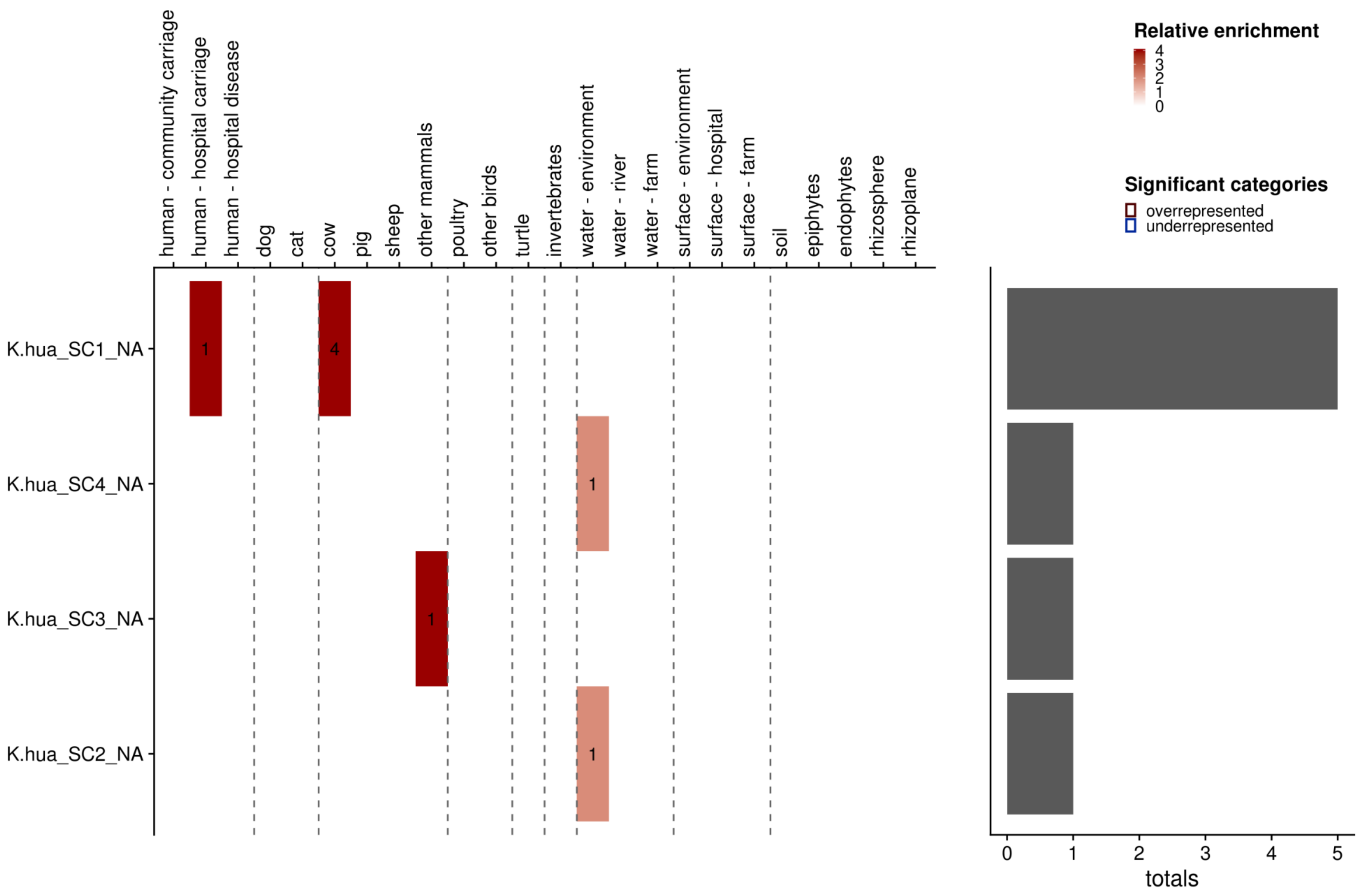

Fig S16 Distribution of *K. ornithinolytica* lineages across different sources

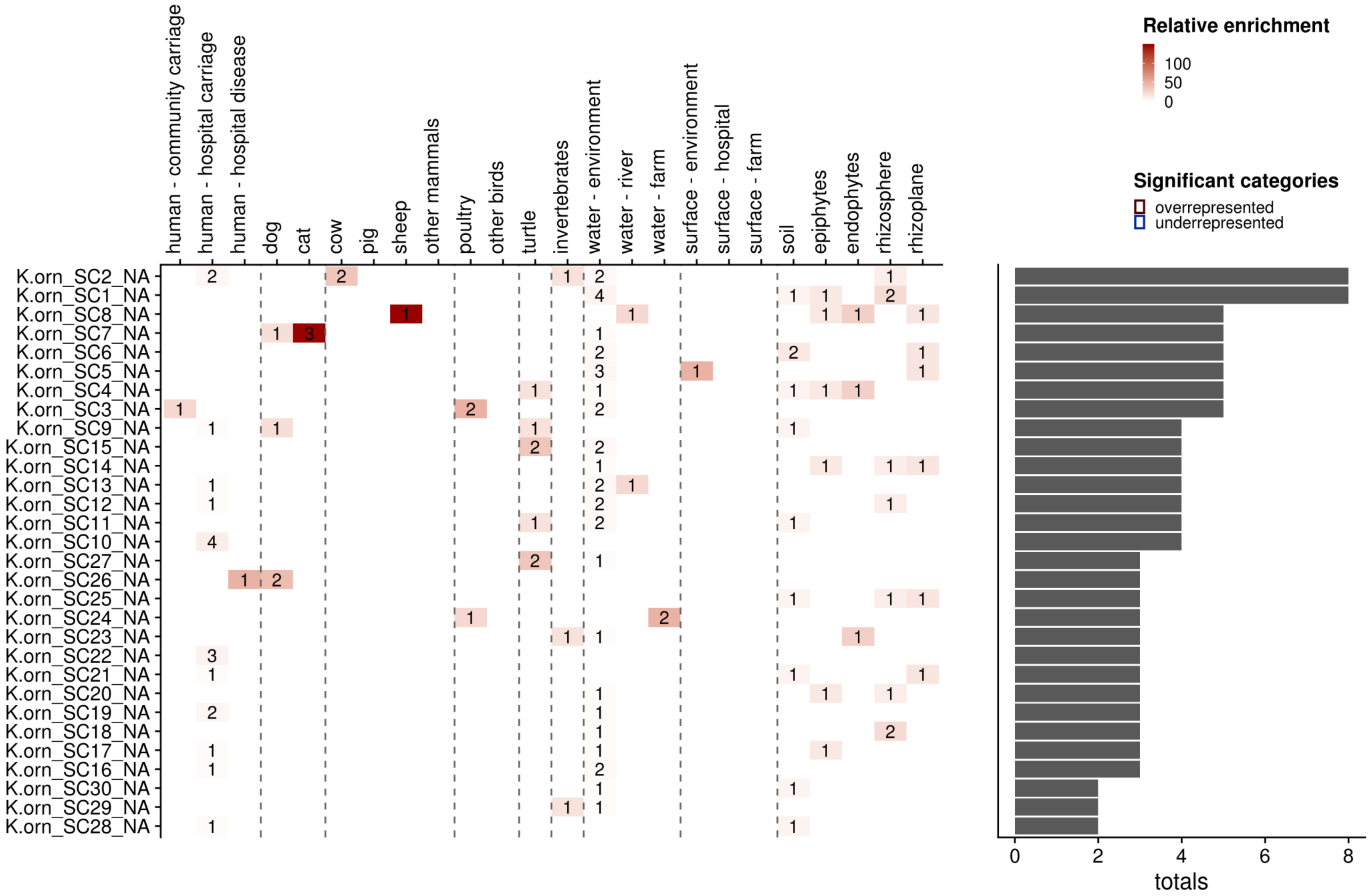

Fig S17 Distribution of *K. planticola* lineages across different sources

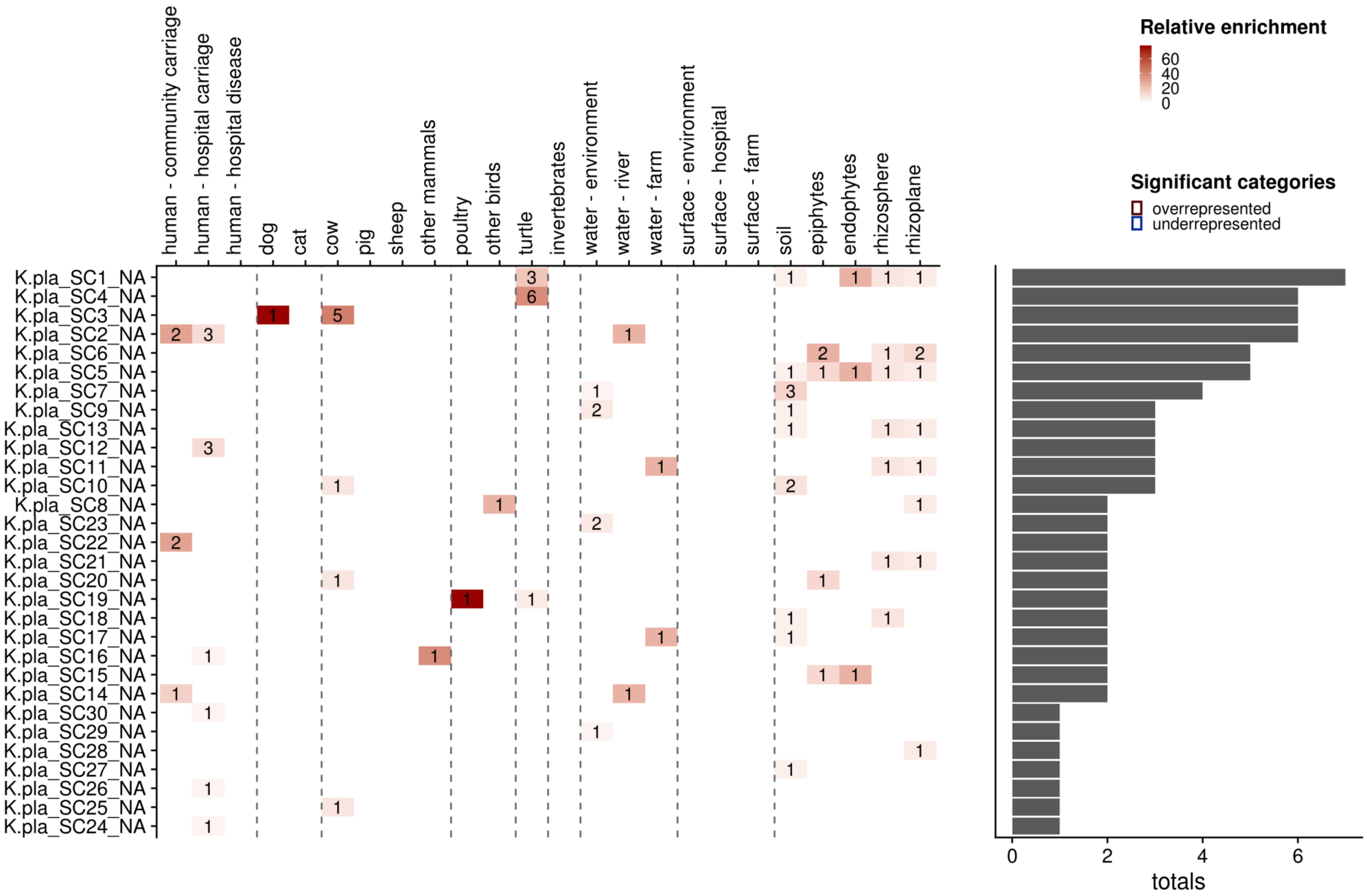

Fig S18 Distribution of *K. terrigena* lineages across different sources

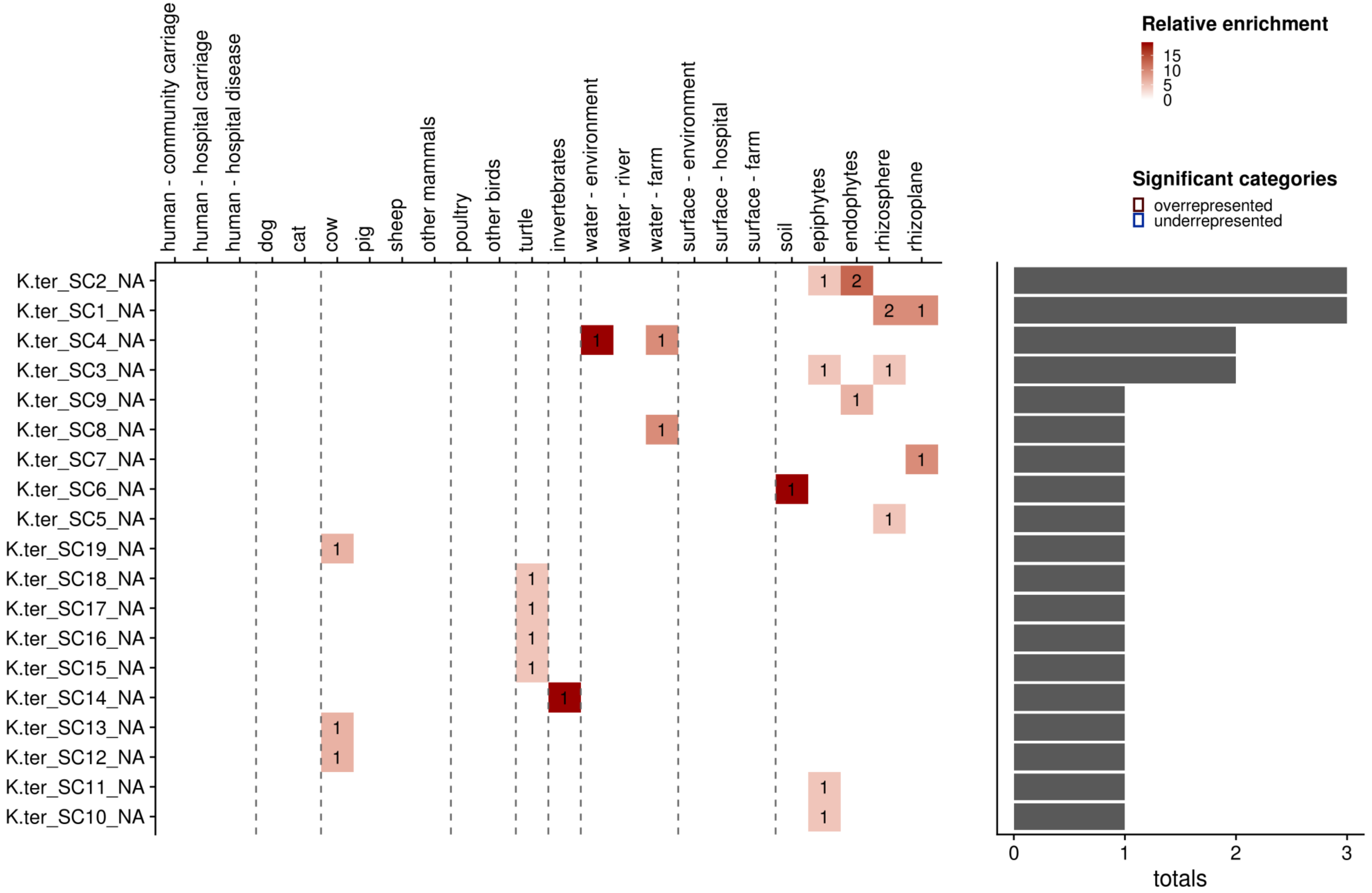

Fig S19 Distribution of *K. quasiterrigena* lineages across different sources

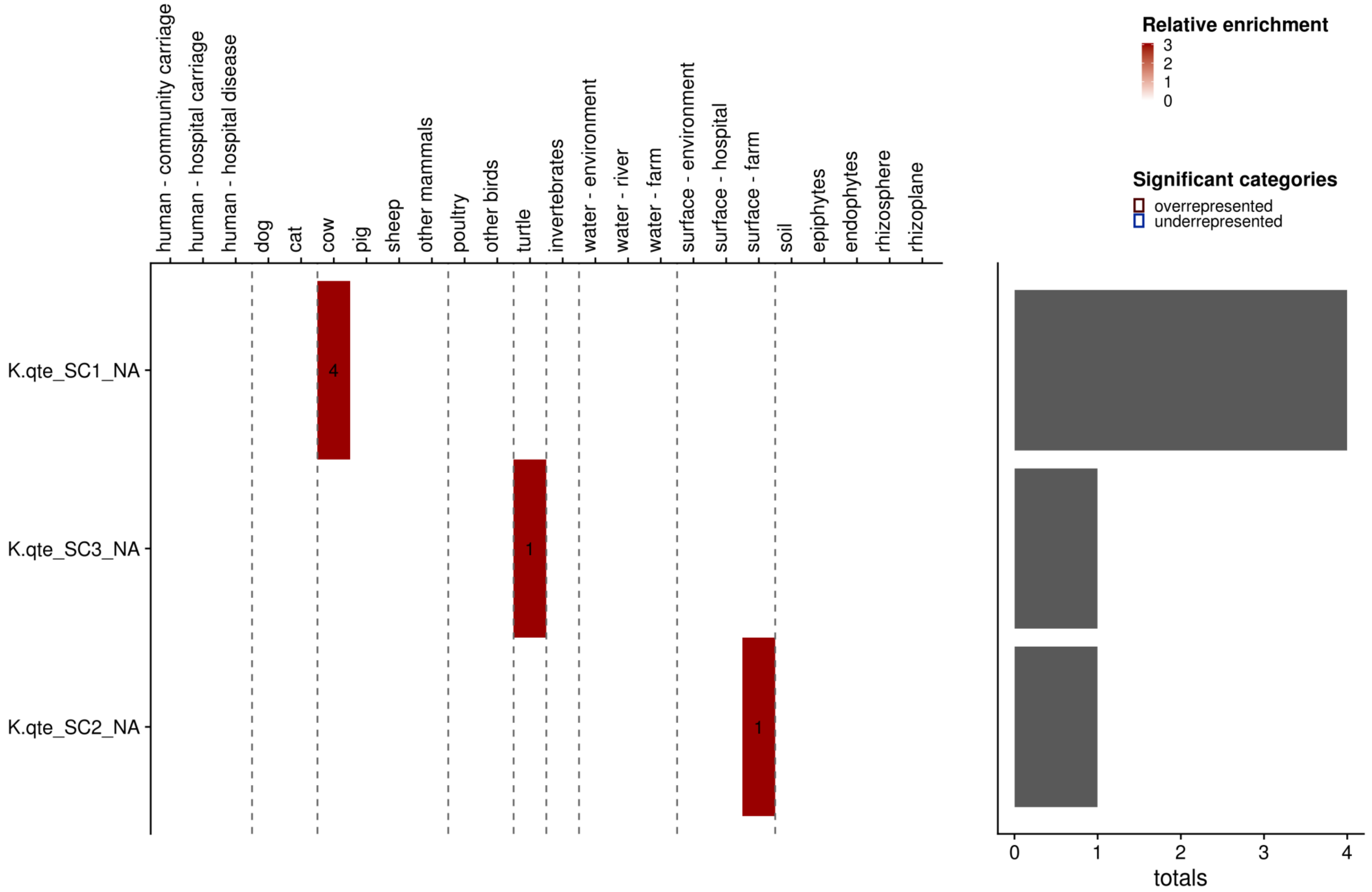

Fig S20 Distribution of *K. aerogenes* lineages across different sources

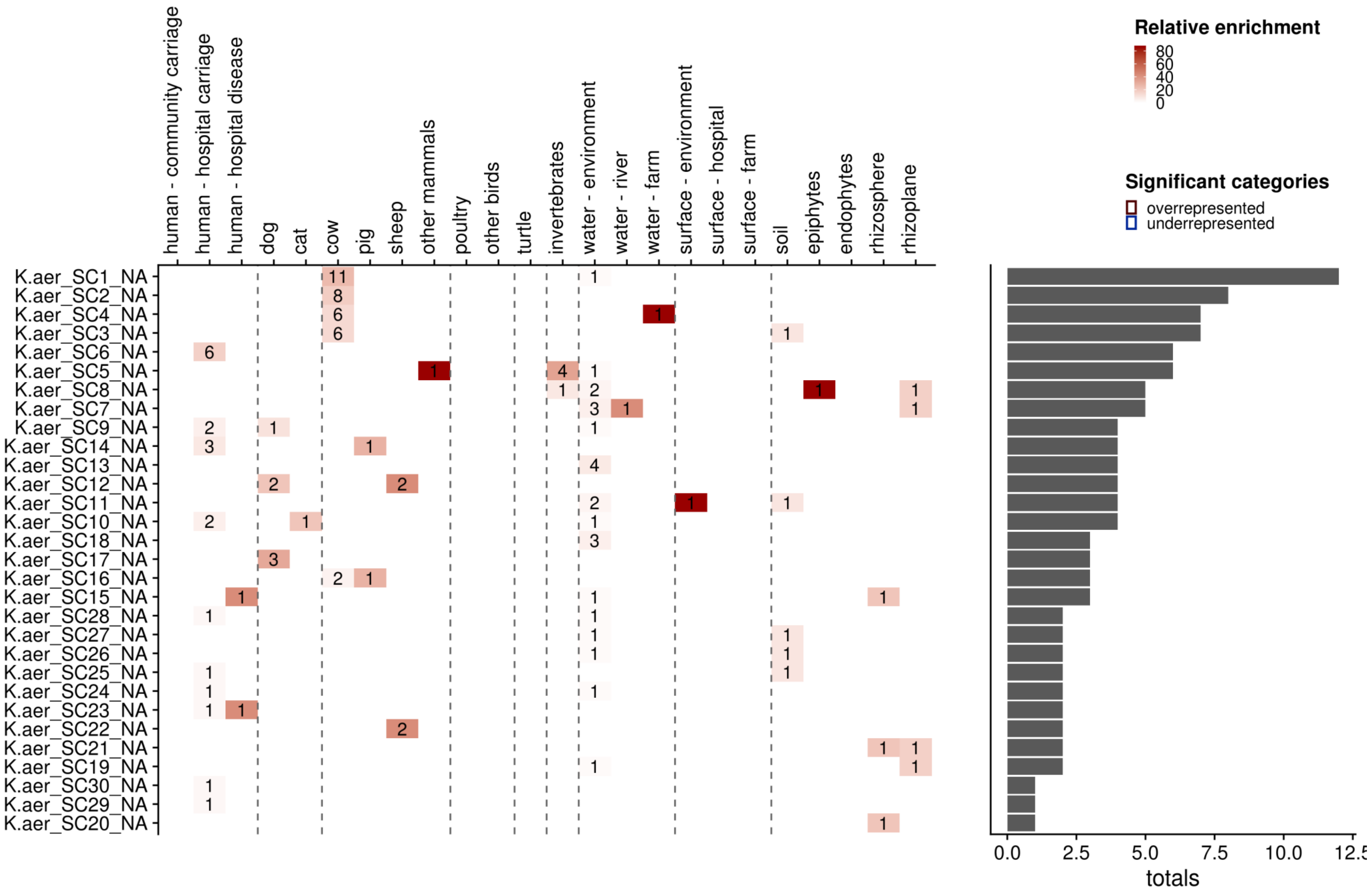

Figure S21

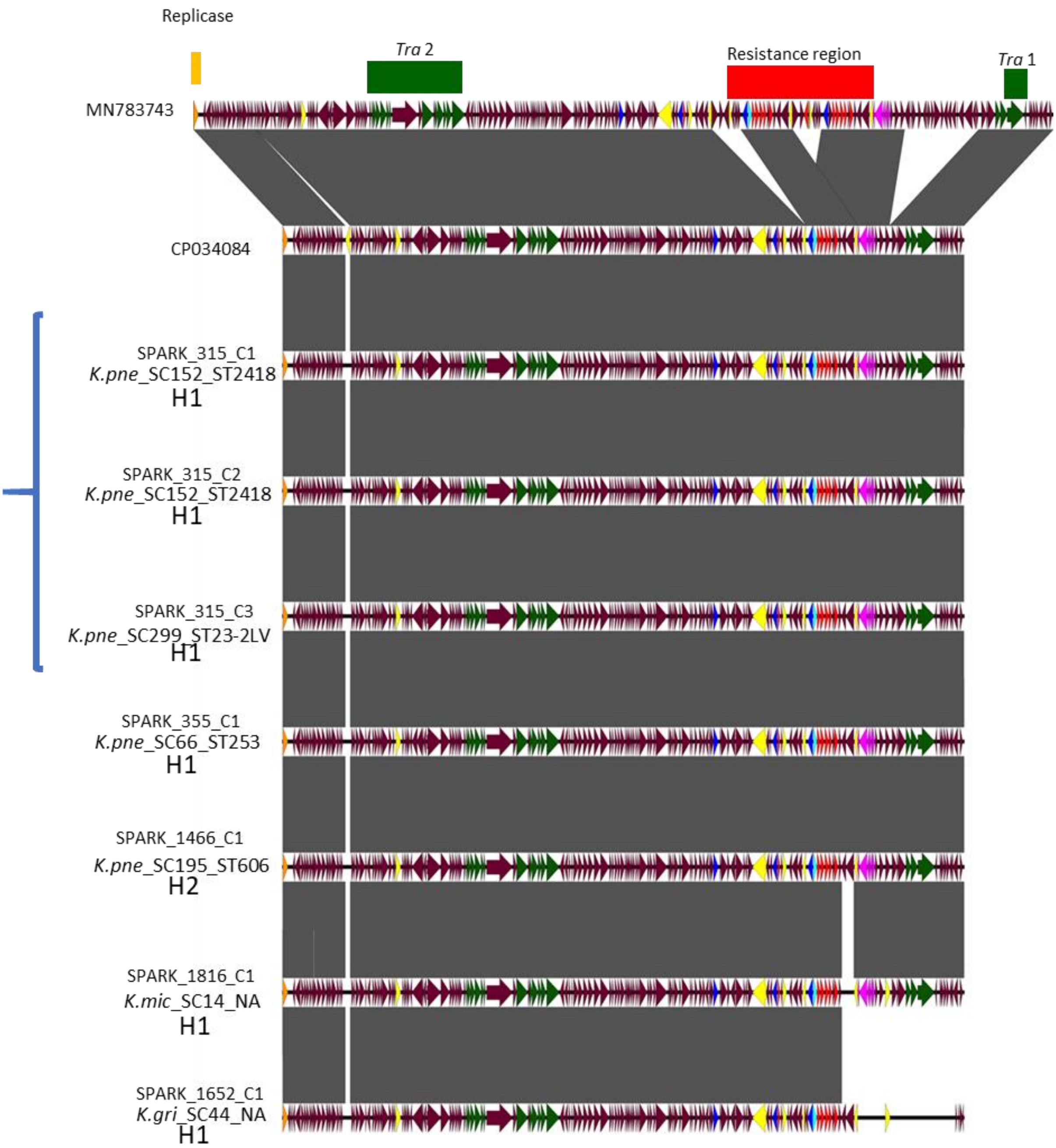


**Figure S21.** Alignment of the MN783743 and CP034084 reference plasmids with the contigs identified in our data as carrying the *bla*$_{VIM}$ gene in IncA/C plasmids. Arrows represent the *bla*VIM gene (turquoise), conjugal transfer system (green), resistance genes (red), mobile elements (yellow), hypothetical proteins (plum), replication protein *repA* (orange), integrases (dark blue) and the mercury resistance operon (pink).

The three bracketed SPARK_315 SCAI carriage isolates were obtained from the same faecal sample from an outpatient in Hospital 1 (H1). Two of these (SPARK_315_C1 and SPARK_315_C2) were members of the same clonal lineage (SC152_ST2418), whilst SPARK_315_C3 was an unrelated isolates related to the hyperviruent lineage ST23 (SC299_ST23-2LV). The plasmids are however essentially identical within all three isolates (which is identical to CP034084), indicating within-patient transfer. SPARK_355_C1 is a diagnostic carriage sample from a rectal swab from a different inpatient in Hospital 1 corresponding to lineage *K.pne*_SC66_ST253 and also contains the same VIM plasmid. SPARK_1466_C1 also contains a very similar plasmid. This isolate corresponds to a different lineage (*K.pne*_SC195_ST606) and is a diagnostic isolate from a urine sample from inpatient in Hospital 2. SPARK_1816_C1 is a K.mic diagnostic isolate (SC14) from a rectal swab from inpatient in Hospital 1 and contains also contains this plasmid. Finally, SPARK_1652_C1 is a K. gri diagnostic isolate (SC44), also from a urine sample from an outpatient in hospital 1. This isolate contains the same plasmid, but with a deletion near the resistance cassette.

Figure S22

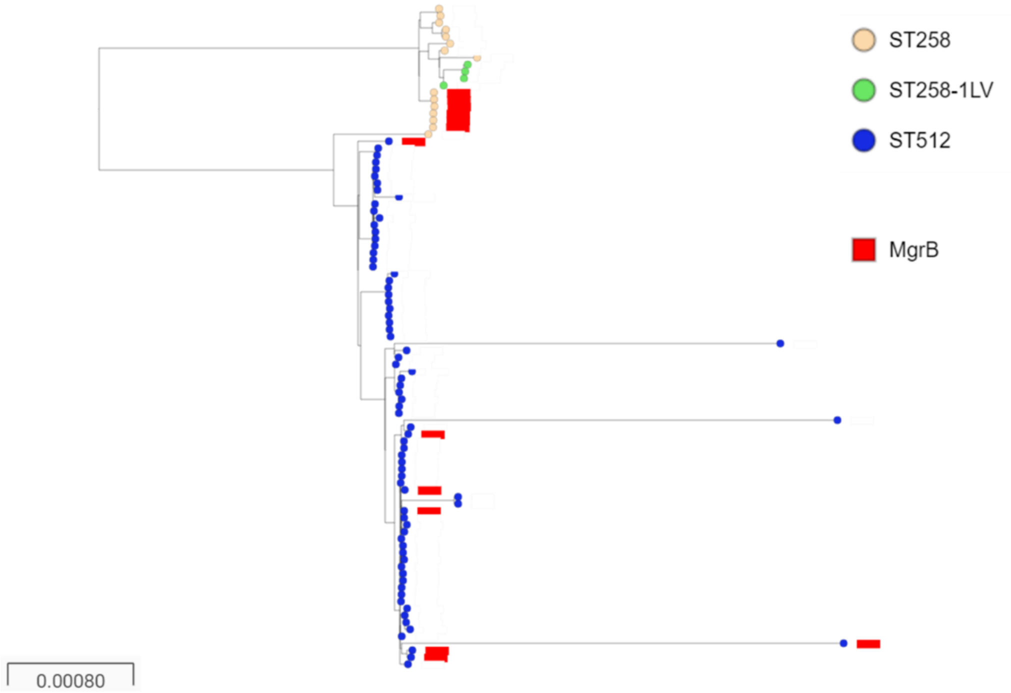


**Figure S22.** The presence of the mutation in *mgr*B conferring colistin resistance in *K. pne* clone ST258/512. The tree was constructed using RaXML

Figure S23

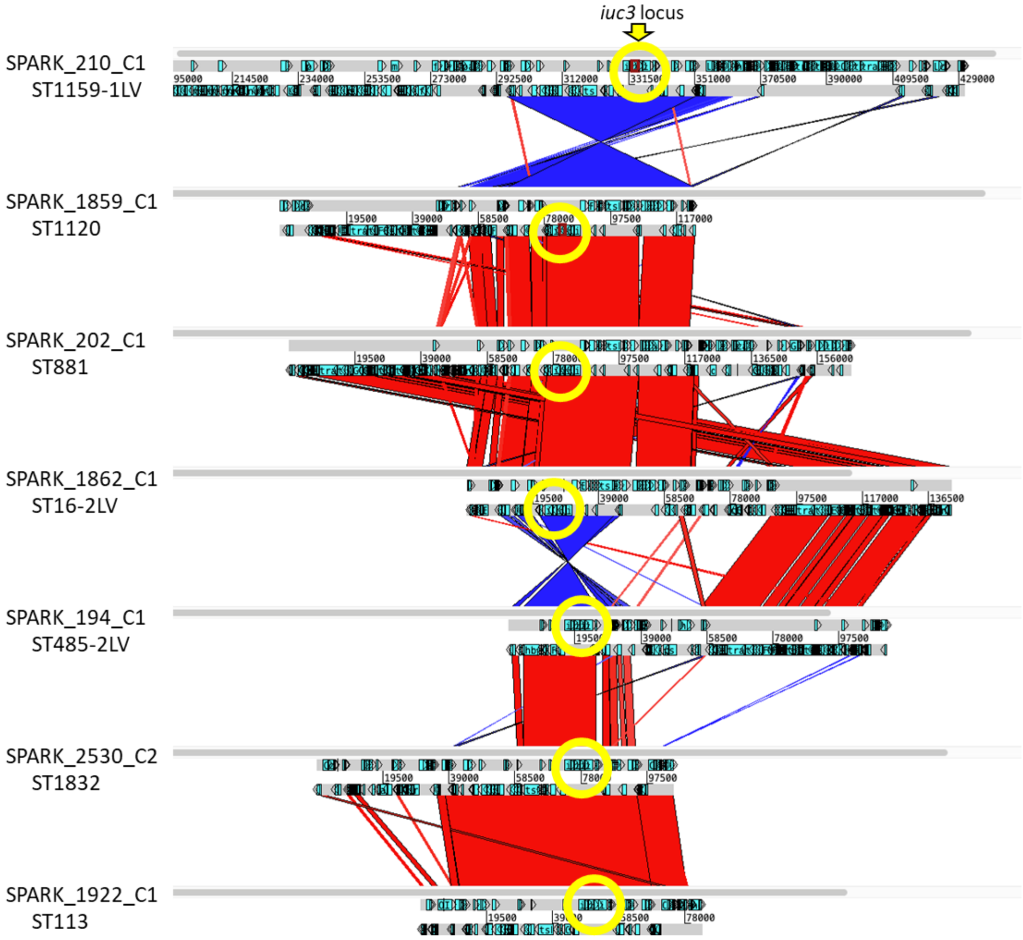


**Figure S23.** Comparison of short-read contigs from *K. pne* isolates from pigs harbouring *iuc3* in this study. The ST and position of *iuc3* are shown. Contigs were compared with the Artemis Comparison Tool (ACT); sections of similarity of >500bp are shown.

Figure S24

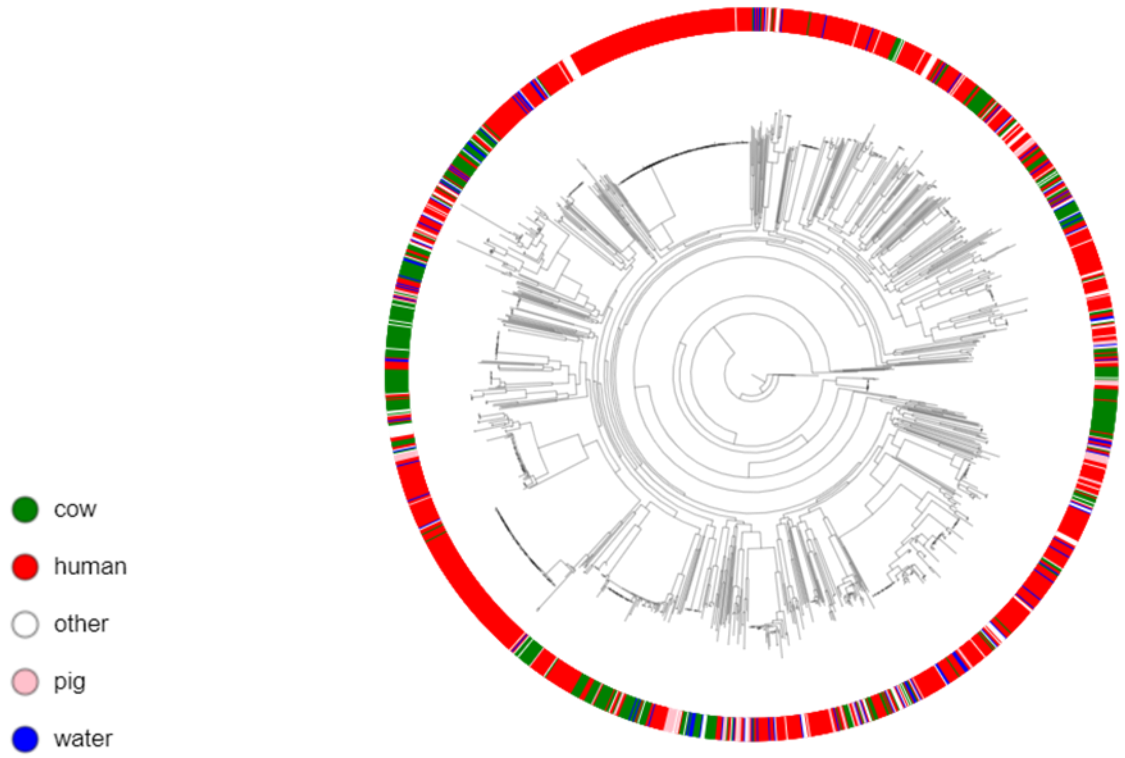


**Figure S24.** Human (red) and cow (green) isolates of *K. pne* are non-randomly distributed across the phylogenetic tree (constructed using RaXML)

**Figure S25**

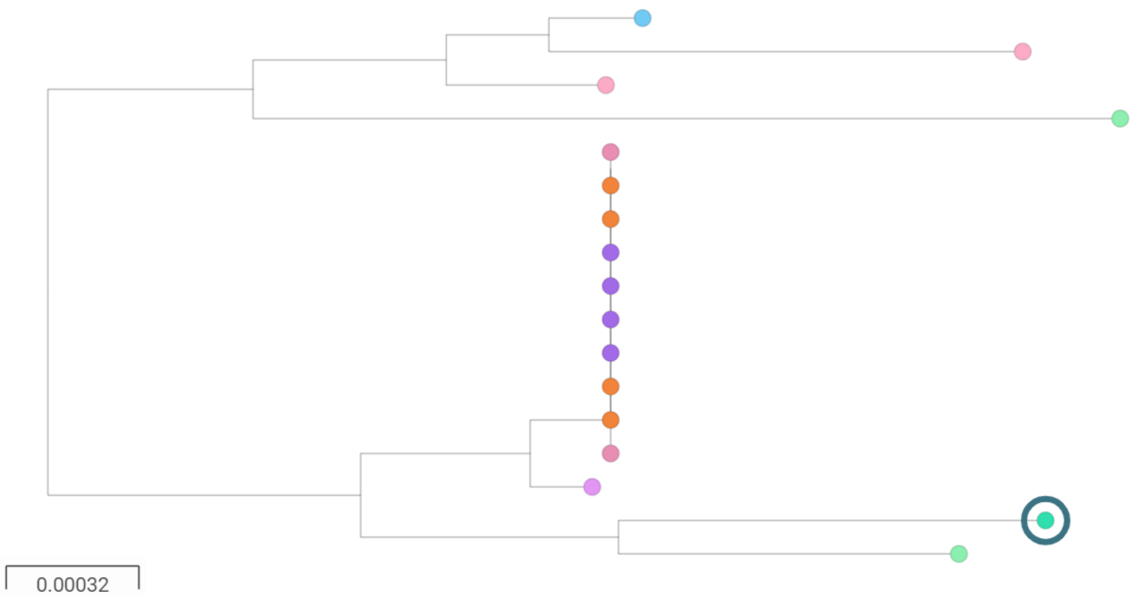


**Figure S25.** *K.gri* SC1 isolates. Most isolates are associated with invertebrates, a single isolate from a hospital outpatient is ringed in the tree

Figure S26

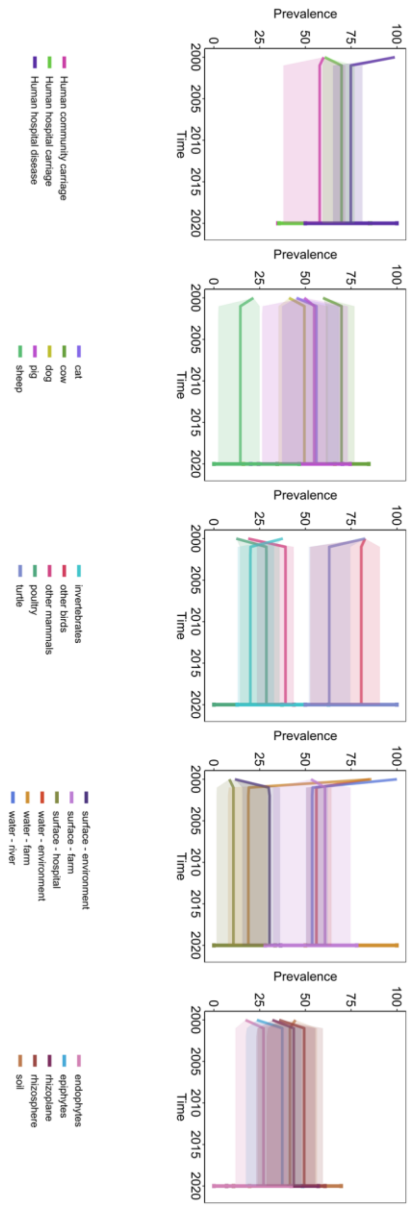


**Figure S26**. All 24 nodes in the connected system and their uncertainty ranges when fit to prevalence estimates. All simulations achieve steady state in 20 years and are between the upper and lower bounds for each node.

Figure S27

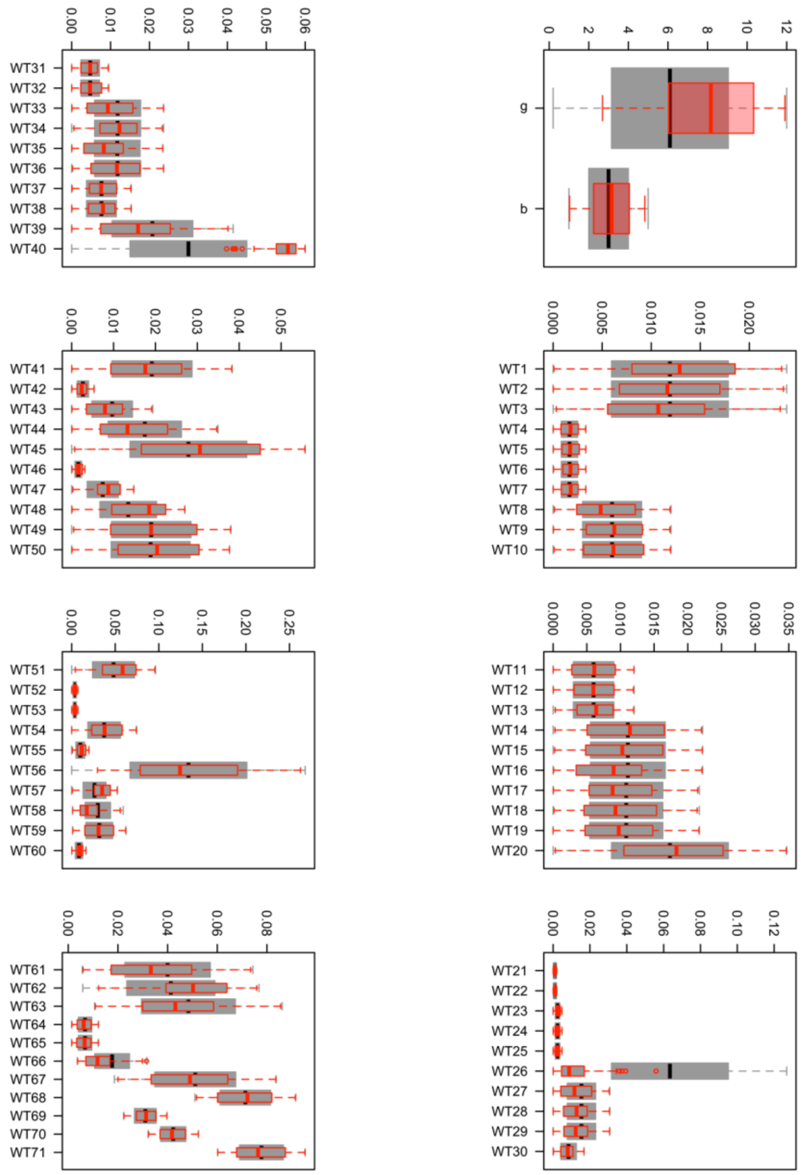


**Figure S27**. The range for each Latin Hypercube sample (grey) and the distribution of the best 135 parameter sets (red) which replicate the transmission network and prevalence data. Transmission weight priors (WT1, …, WT71; Table S1) are calculated as the number of transmission events between nodes, divided by the total samples taken within the 'from' node with 95% credible interval.

**Figure S28**

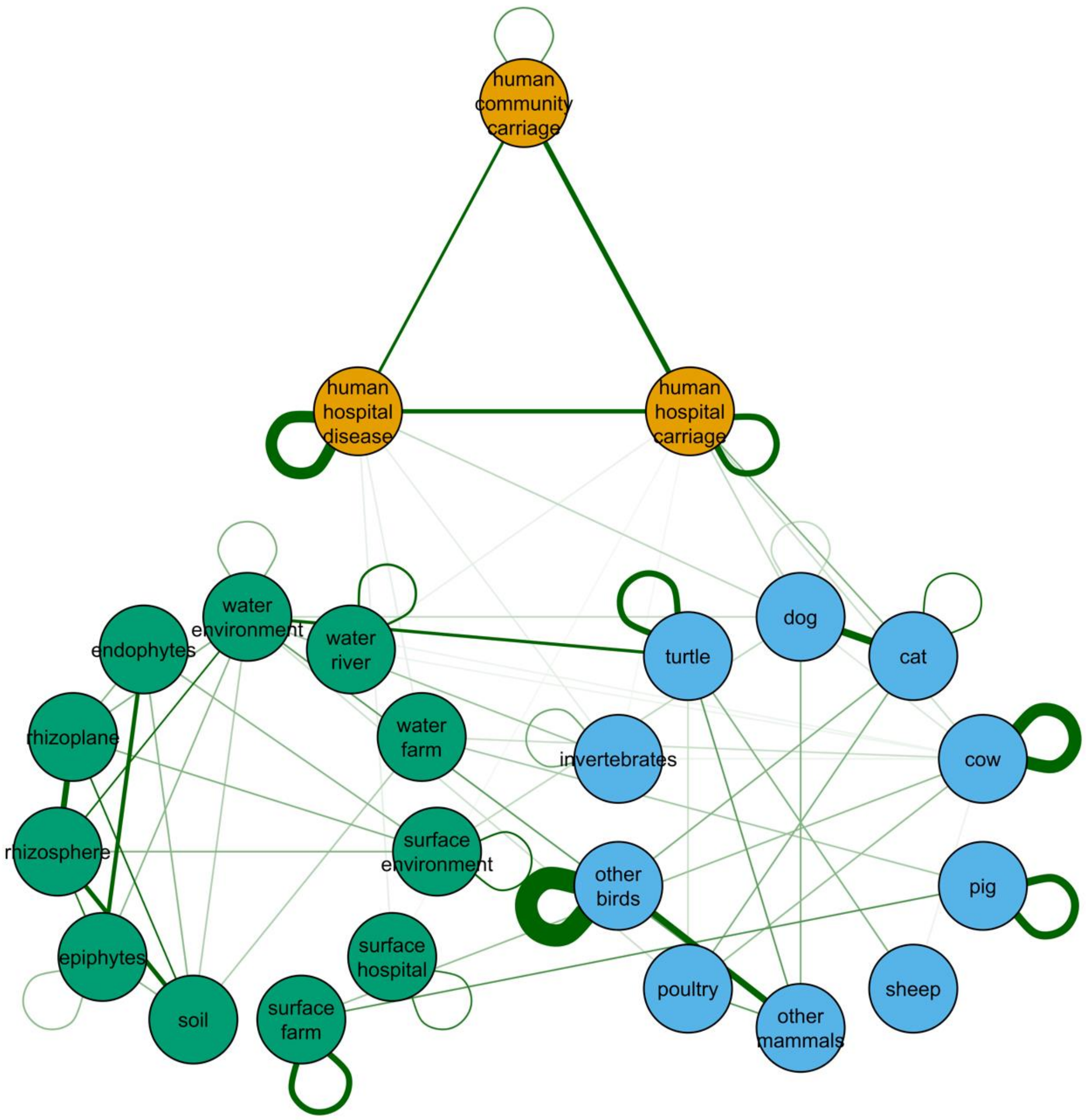


**Figure S28**. All 24 nodes and their median relative transmission weight in the connected system of humans (orange), animals (blue) and environment (green). Thicker links between nodes indicate higher relative transmission. We sample from the full range in order to explore uncertainty.

Figure S29

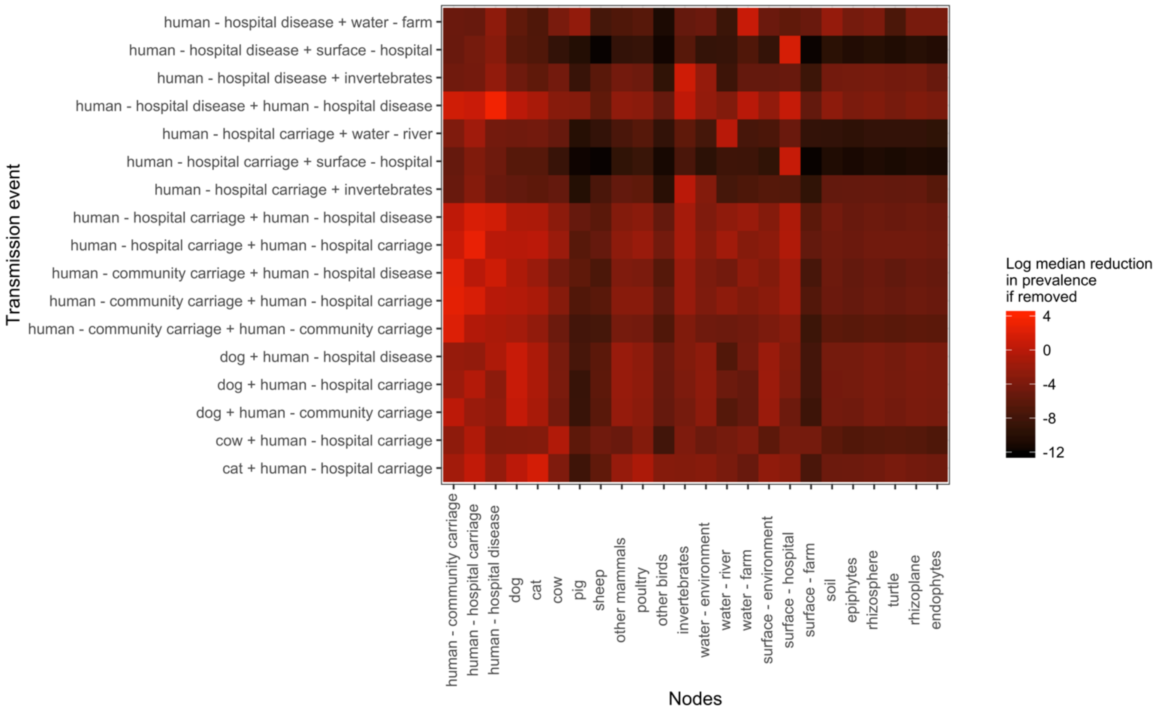


**Figure S29a**. The median projected impact (reductions in prevalence) over 20 years of removing each transmission event relating to **humans** (y axis) on each of the 24 nodes (x axis)

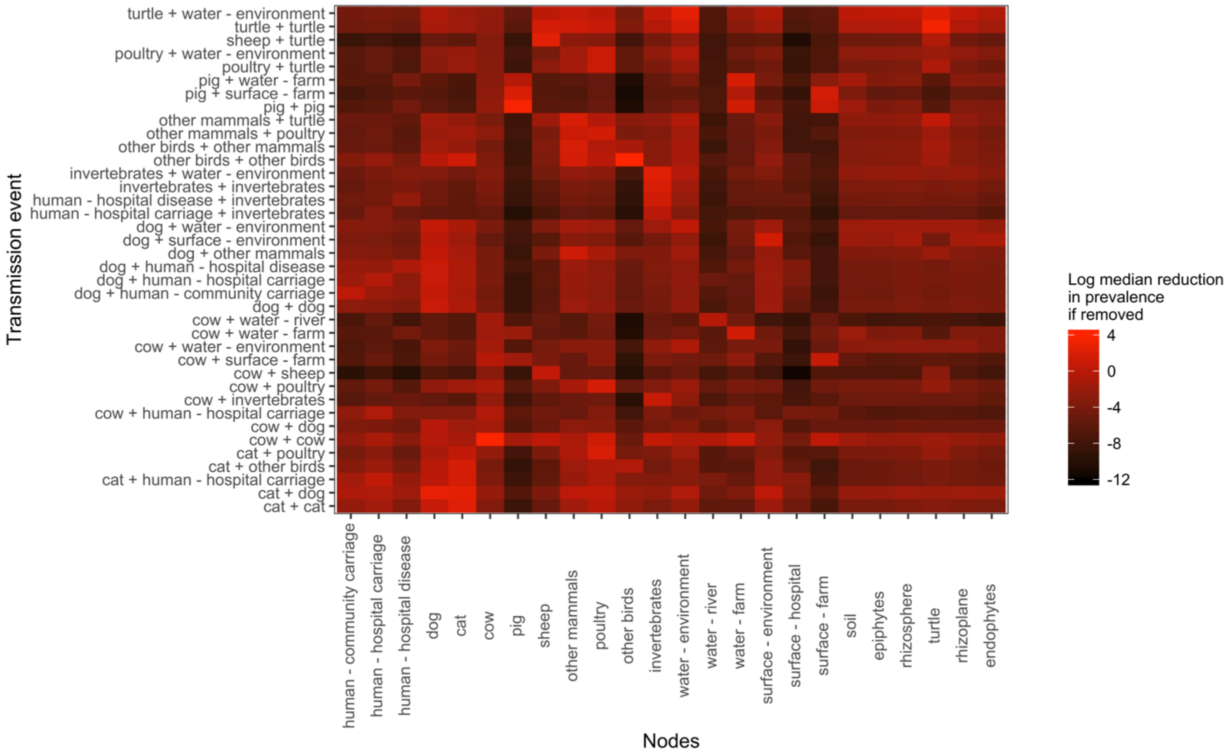

**Figure S29b.** The median projected impact (reductions in prevalence) over 20 years of removing each transmission event relating to **animals** (y axis) on each of the 24 nodes (x axis)

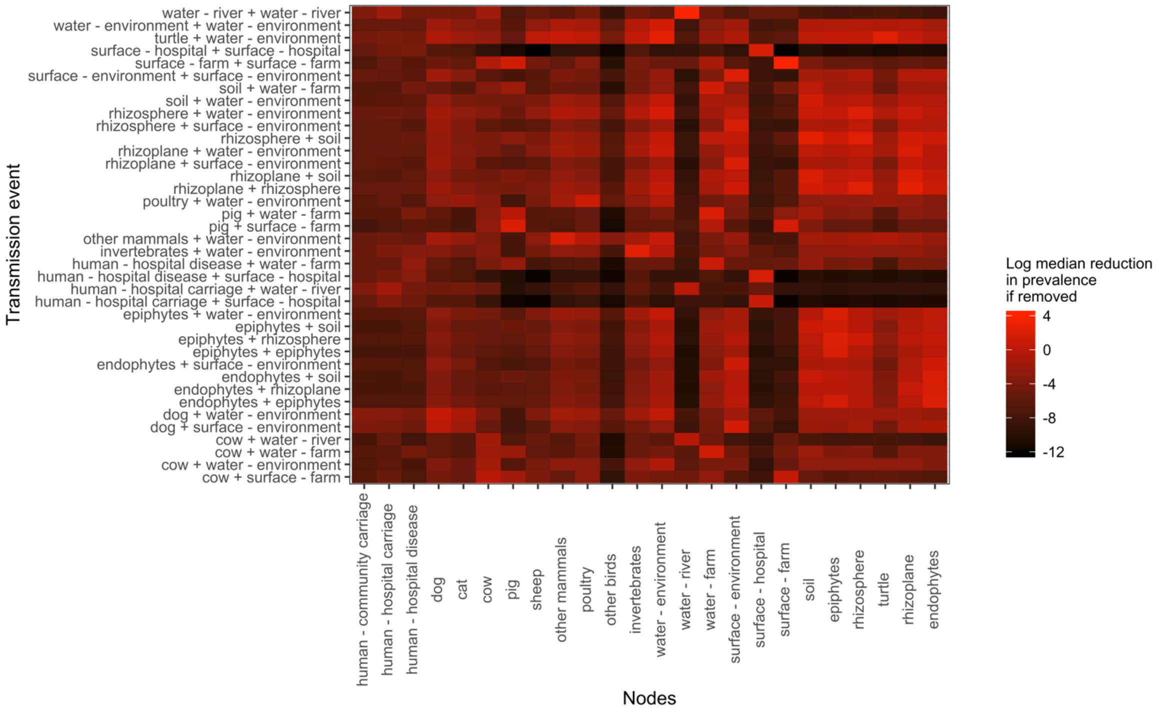


**Figure S29c**. The median projected impact (reductions in prevalence) over 20 years of removing each transmission event relating to the **environment** (y axis) on each of the 24 nodes (x axis)
