## Supplementary notes for "One Health or Three? Transmission modelling of *Klebsiella* isolates reveals ecological barriers to transmission between humans, animals and the environment"

**Supplementary note 1*:* Newly described species and novel lineages**.

**Recently described species:**
Our WGS data includes isolates of two newly described species *K. spallanzani* (*K.spa*; n=10) and *K. pasteurii* (*K.pas*; n=27) that were recovered for the first time during this study (Merla *et al*), and 8 isolates of the recently described *K. huaxensis (K.hua)* that has only previously only been recovered from a urine sample from China[1]. All these species belong to the *K.oxy*SPEC (Fig S1). *K.pas* is a sister-species to *K.gri*, and the 27 *K.pas* isolates were recovered form a wide range of human, animal and environmental niches. Two unrelated *K.pas* strains (SPARK_1778_C1, and SPARK_2511_C1) were recovered from urine samples from inpatients of two different hospitals; these are the first examples of this species being associated with a urinary tract infection. *K.pas* Isolate SPARK_1778_C1 harboured the *qnr*-S1 gene and was phenotypically resistant to ciprofloxacin and levofloxacin.

*K.spa* and *K.hua* represent the most diverged branch of the *Koxy*SPEC (Fig S1). Isolates of both species were recovered from diverse environmental and animal settings. A single isolate (775_C1) of *K.spa* was recovered from the urine of a hospital outpatient and showed phenotypic resistance to fosfomycin, as previously described in Merla et al[2]. The single isolate of *K.hua* previously reported from China[1] was also recovered from urine of an ICU patient, but of the 8 strains of this species in our data, none were recovered from human disease; instead, 4 were recovered from cows, 2 from water, 1 from a horse and one from hospital carriage.  There was no notable genotypic or phenotypic resistance in the *K.hua* isolates.

**Novel Lineages:**
The two isolates SPARK_361_C1 and SPARK_2596_C1 correspond to a novel lineage of ambiguous species status. Both isolates were from hospital carriage but were not epidemiologically linked and contain no notable resistance or virulence factors. These two isolates are over 1% diverged from each other, which is towards the upper end of typical divergence within a single species in our dataset. They fall at an intermediate position between K.pas (>3% divergence) and K. gri (>2.5% divergence). Thus, they are not sufficiently diverged from either of these species to warrant separate species status, yet it is unclear as to which of these species they should be assigned.

We also recovered six isolates of a novel lineage to which we have applied the label *K. quasiterrigena* (*K.qte*). This group clusters within the *Korn*SPEC species complex (Fig S2), and is >95% diverged in terms of ANI  from the most closely related species *K.ter*  (not shown). Four of the *K.qte* isolates belong to a single bovine clone (K.qte_SC1), the other two isolates were isolated from the farm environment, and one from a turtle. There is no notable genotypic or phenotypic resistance in the *K.qte* isolates.


1. Hu, Y., Wei, L., Feng, Y., Xie, Y. & Zong, Z. Klebsiella huaxiensis sp. nov., recovered from human urine. Int. J. Syst. Evol. Microbiol. 69, 333–336 (2019)
2. Merla, C. et al. Description of Klebsiella spallanzanii sp. nov. and of Klebsiella pasteurii sp. nov. Front. Microbiol. 10, 2360 (2019)

**Supplementary Note 2 : Diagnostic isolates**

In addition to the 2795 SCAI sequenced isolates, we sequenced an additional 687 clinical *Klebsiella* isolates recovered as part of ongoing surveillance programs within the Pavia catchment area. The full metadata for these strains is available via the microreact project. 600/687 (87.3%) of the isolates were from human clinical cases, 513(89%) from hospital disease, 76 (11%) from hospital carriage and 11 (0.1%) from companion animals. The human-derived isolates were recovered from four hospitals with the majority being from a single hospital (n= 484; 70%). The majority of the 600 clinical isolates were recovered from urine (n=421; 70.1%), the other sample types being blood (n=74; 12.3%), rectal swab (n=70; 11.6%), bronchial (n=45; ), sputum (n=20; 7.5%) and other minor sources. 578 of all 687 isolates were assigned by WGS as *K.pne* (84.1%), with the other isolates being, *K.oxy* (n=40), *K.var* (n=23), *K.mic* (n=18), *K.qpq* (n=7), *K.qps* (n=5), *K.gri* (n=6), *K.aer* (n=3) and *K.orn* (n=3), *K.pas* (n=2), *K.qva* (n=1), *K.spa* (n=1). This confirms the dominance of *K.pne* as a cause of human infection, with the recognised opportunistic pathogen *K.oxy* being the second most common source. However, we note that *K.mic* is the fourth most common species from the hospital disease sample, and is recovered almost as commonly as the more recognized opportunistic pathogen *K.var* in third place. The 18 isolates of *K.mic* recovered from human disease (including one SCAI isolate) were not clonally or epidemiologically related, and were recovered from a variety of sample types.

**Supplementary Note 3: A mathematical model for hierarchy of transmission networks**

**Model fitting**

We fitted the model on a total of 1000000 parameter sets, and each simulation was either accepted or rejected based on how well the prevalence for each node reflected the real data. The majority of final prevalence values were able to be fit, but for one node, *water – farm,* the transmission events did not align well with the final prevalence (31/36 positive), therefore, we were unable to replicate the prevalence within this single node. However, 23 from the total 24 nodes were fit, thus ensuring the transmission network adequately reflected the remaining prevalence found in each node in 2020. From 1 000 000 Latin Hypercube samples, 135 sets of parameters reflected the transmission data and prevalence. These 135 runs are used as the final fitted set of parameters and are equilibria states shown in Figure S26. These 135 runs represent the best guess for each parameter, while incorporating uncertainty in the final parameter range (shown in Figure S27).

**Intervention Analysis**

Each of the 71 transmission events (Table S7) were removed one-by-one to project the impact on the transmission dynamic system over a time period of 20 years. Figures S29a (humans), S29b (animals) and S29c (environment) show heat maps to illustrate the median change in each node when removing each transmission event from the transmission network. This gives an indication of the 'connectedness' of all nodes and shows the impact each transmission event has further into the future (2020 – 2040). The darker the colour, the less impact each transmission event has on each node, with lighter shades of red indicating higher impact by removing those transmission events (on log scale).

Figure S30 shows the impact of removing each node on the percentage median reduction in the prevalence of *human – hospital disease* over 20 years. This illustrates the total projected impact of each node (and therefore all of its connected transmission links) on *human – hospital disease.*
